# Helix-8 and Carboxyl Terminal Tail Regulation of Proteinase Activated Receptor-4 (PAR4) signalling

**DOI:** 10.1101/2025.02.10.637468

**Authors:** Pierre E. Thibeault, Amr A.K. Mousa, Victor M. Mirka, Rithwik Ramachandran

## Abstract

The 8th-helix (H8) and carboxyl terminal tail (CT) of GPCRs are crucial for interactions with intracellular signaling molecules and regulatory proteins. We investigated how H8 and CT residues influence signaling in PAR4, a tethered-ligand activated GPCR. Our analysis revealed that PAR4 activation by thrombin or AYPGKF-NH2 stimulates β-arrestin-1/-2 recruitment and activates multiple Gα proteins (Gαq, Gα11, Gαz, Gα15, Gα12, Gα13, Gαi1-3, GαoA/B). Mutation of the H8 sequence (R^352^AGLFQRS^359^) significantly reduced G protein activation and β-arrestin recruitment, with residues Leu^355^-Glu^357^ being particularly important. Mutations of specific lysine residues in H8 and CT also impaired signaling. Additionally, we identified a crucial TM7-H8 interaction and found that CT phosphorylation sites regulate β-arrestin recruitment. Finally, using AlphaFold3 we predicted interactions between PAR4, β-arrestins, and G proteins, revealing novel receptor regulatory sites in the intracellular loops, transmembrane domains, H8 and CT.

## Introduction

G protein-coupled receptors (GPCRs) are the largest family of cell membrane signalling proteins expressed in eukaryotic cells^1,2^. These receptors convert extracellular ligand binding into intracellular signalling to mediate cellular affects. Ligand binding activates the GPCR through a series of structural reorganizations that allow for effectors, such as G proteins, to bind and become activated. In addition to the intracellular loops of GPCRs, the H8 and CT also play important roles in effector interaction.

Proteinase activated receptors (PARs) are a family of four (PAR1-4) rhodopsin-like (Class A) GPCRs, which are activated by a host of proteolytic enzymes from coagulation cascade-, immune cell-, and pathogen-derived sources^3–5^. PARs are unique compared to other Class A GPCR counterparts in their mechanism of activation which involves proteolysis of the receptor N-terminus to reveal a novel sequence, often termed the tethered ligand, that is capable of activating the receptor through intramolecular binding^3^. Alternatively, subtype-selective PAR activation can be achieved in the absence of proteolytic cleavage, through application of synthetic tethered ligand-mimetic peptides^6–9^.

PAR4 performs important functions in regulating thrombosis and hemostasis through its role as a platelet thrombin receptor^3,10–12^. In addition to thrombin, PAR4 is also activated by other enzymes including trypsin and neutrophil Cathepsin-G^5^. Proteolytic cleavage of PAR4 by thrombin unmasks the tethered ligand sequence, G^48^YPGQV.., which binds intramolecularly to the receptor, leading to its activation. Alternatively, tethered ligand-mimicking peptides, such as AYPGKF-NH_2_, can activate PAR4 without proteolytic cleavage^7,13^. PAR-mediated platelet activation downstream of thrombin release at the site of vessel injury is well-described, and as such, these receptors have been the target for anti-platelet therapeutic approaches^10,11,14,15^.

When activated with either the thrombin cleavage-revealed tethered ligand, G^48^YPGQV^53^…, or tethered ligand-mimicking peptide, AYPGKF-NH_2_, PAR4 engages Gα_q/11_, Gα_12/13_, and Gα_i/o_ G protein subtypes as well as β-arrestins^7,13,16,17^, although PAR4 cupling to Gα_i_ independent of crosstalk with purinergic receptors has been questioned in certain tissue contexts^7,16,18–20^. Previously, we demonstrated key differences in CT regulation of β-arrestin-1/-2 recruitment and G protein mediated signaling between peptide and proteolytic activation of the related PAR2 receptor^21^. In contrast to PAR2 and unlike many other Class A GPCRs, PAR4 lacks certain conserved residues and motifs implicated in effector interaction and signalling, such as a H8 cysteine residue and CT clusters of serine/threonine residues. Thus, the molecular mechanisms that underlie PAR4 signalling, and signal regulation are of interest – both for understanding the mechanisms of effector interactions with PAR4 as well as for the development of targeted therapeutic strategies. Further, probing differential contributions of residues to signalling and regulation downstream of either peptide activation or proteolytic cleavage provides insight into how these two different modes of activation may differentially regulate these interactions.

In this study, we use Alphafold modelling and mutagenesis to determine H8 and CT residues and motifs important for thrombin or AYPGKF-NH_2_ activated PAR4 interaction with signaling effectors. We find that residues in both the H8 and CT differentially impact PAR4 interaction with G proteins and β-arrestins. Together our findings provide novel insights into molecular mechanisms regulating PAR4 functions and may aid in the development of novel therapeutics targeting this receptor.

## Results

### G protein activation profile of PAR4 in response to activation with the tethered-ligand mimicking peptide, (AYPGKF-NH_2_) or thrombin

Our understanding of the G protein-coupling profile of PAR4 is largely based on second messenger signaling responses. To gain an understanding of G protein subtype selectivity and agonist-dependent differences, we performed TRUPATH assays following activation of PAR4 with the thrombin revealed tethered-ligand GYPGQV…, or the PAR4 tethered ligand-mimicking peptide AYPGKF-NH_2_ to assess the G protein subtypes that are activated. TRUPATH is a BRETII assay wherein 14 Gα-Rluc8 proteins are co-transfected with untagged Gβ, and Gγ-GFP2^22^. Decrease in BRETII signal signals a change in conformation or dissociation of the βγ complex from the Gα protein – indicative of G protein activation^22^. We transfected HEK-293 or CRISPR/Cas9 PAR1-knockout (PAR1-KO) HEK-293 cells ^17^ with PAR4 and the various TRUPATH Gα-β-γ constructs and monitored responses to AYPGKF-NH_2_ and thrombin. We observed that AYPGKF-NH_2_ activation of PAR4 stimulated Gα_q_, Gα_11_, Gα_z_, Gα_15_, Gα_12_, Gα_13_, Gα_i1_, Gα_i2_, Gα_i3_, Gα_oA_, Gα_oB_, but not Gα_sS_, Gα_sL_, or Gα_Gustducin_ (Gα_q_ EC_50_ = 18.4 ± 6.1 µM, Gα_11_ EC_50_ = 5.9 ± 2.0 µM, Gα_z_ EC_50_ = 35.1 ± 7.4 µM, Gα_15_ EC_50_ = 2.9 ± 1.9 µM, Gα_12_ EC_50_ = 1.6 ± 0.5 µM, Gα_13_ EC_50_ = 2.8 ± 0.6 µM, Gα_i1_ EC_50_ = 24.5 ± 5.6 µM, Gα_i2_ EC_50_ = 24.4 ± 10.4 µM, Gα_i3_ EC_50_ = 29.3 ± 5.4 µM, Gα_oA_ EC_50_ = 9.7 ± 2.0 µM, Gα_oB_ EC_50_ = 8.5 ± 1.5 µM, Fig. 1). Following activation of PAR4 with thrombin, we observed that PAR4 triggered Gα_q_, Gα_11_, Gα_z_, Gα_15_, Gα_12_, Gα_13_, Gα_i1_, Gα_i3_, Gα_oA_, Gα_oB_ and similarly did not Gα_sS_, Gα_sL_, or Gα_Gustducin_. In contrast to AYPGKF-NH_2_, thrombin activation of PAR4 did not stimulate Gα_i2_ activity (Gα_q_ EC_50_ = 0.27 ± 0.17 Units/mL, Gα_11_ EC_50_ = 0.25 ± 0.01 Units/mL, Gα_z_ EC_50_ = 2.06 ± 0.54 Units/mL, Gα_15_ EC_50_ = 0.27 ± 0.18 Units/mL, Gα_12_ EC_50_ = 0.04 ± 0.01 Units/mL, Gα_13_ EC_50_ = 0.17 ± 0.06 Units/mL, Gα_i1_ EC_50_ = 0.70 ± 0.31 Units/mL, Gα_i3_ EC_50_ = 0.79 ± 0.70 Units/mL, Gα_oA_ EC_50_ = 0.55 ± 0.09 Units/mL, Gα_oB_ EC_50_ = 0.43 ± 0.17 Units/mL; Fig. 2). Thus, PAR4 appears to selectively activate members of the Gα_q/11_, 12/13, and i/o subfamilies, but not Gα_s_ proteins. Further, selectivity within these G protein subfamilies is evident between agonists (Apparent rank order derived from EC_50_ AYPGKF-NH_2_ -Gα_12_, Gα_13_, Gα_15_, Gα_11_, Gα_oB_, Gα_oA_, Gα_q_, Gα_i2_, Gα_i1_, Gα_i3_, Gα_z_; Apparent rank order derived from EC_50_ thrombin – Gα_12_, Gα_13_, Gα_11_, Gα_q_, Gα_15_, Gα_oB_, Gα_oA_, Gα_i1_, Gα_i3_, Gα_z_).

**Fig. 1.**
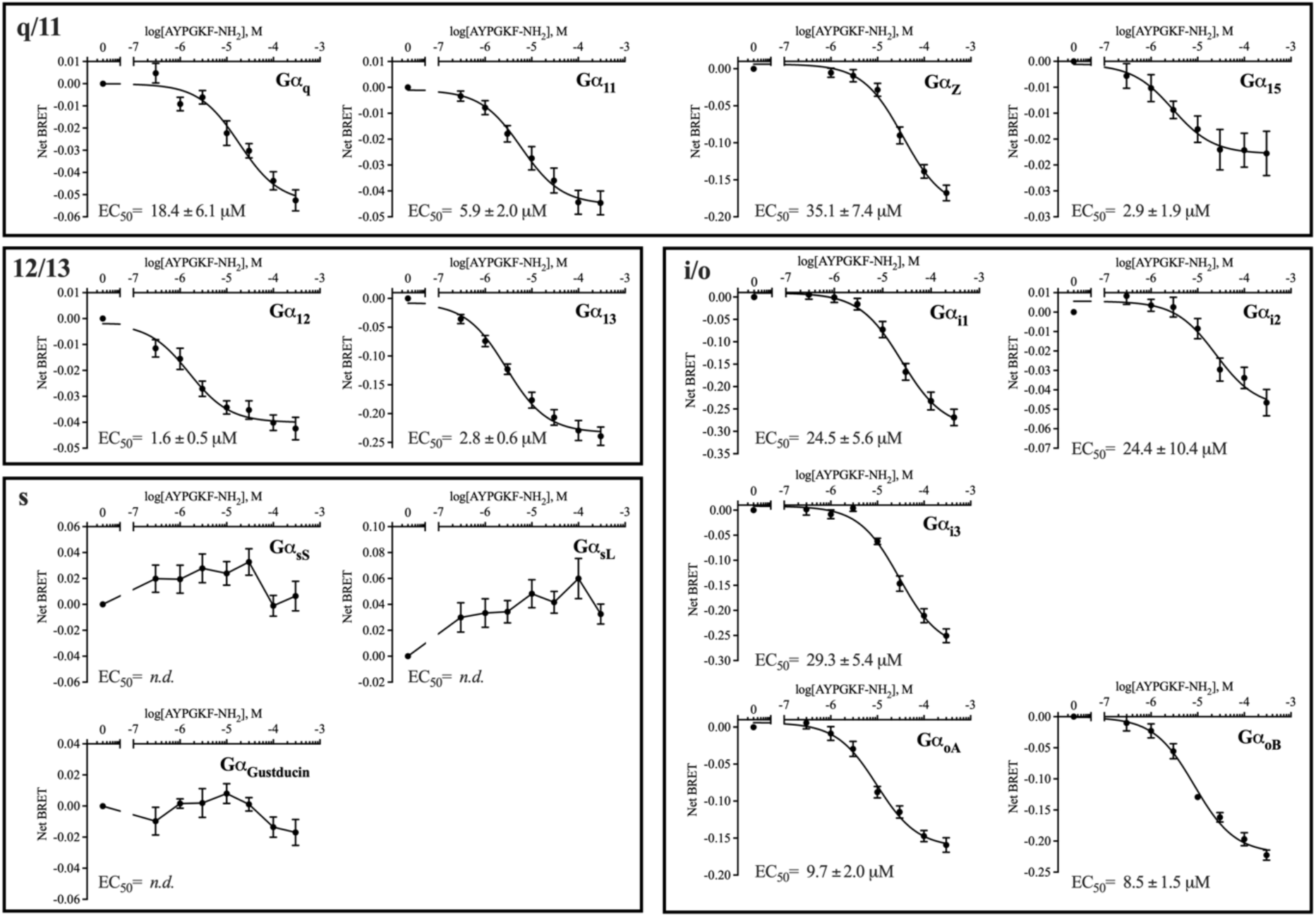
AYPGKF-NH_2_-activated PAR4 stimulates Gα_q/11_, Gα_12/13_, and Gα_i/o_ G protein-subtypes, but not Gα_s_ family subtypes. Assays with HEK-293 cells expressing wild-type PAR-YFP receptor were conducted for each of the TRUPATH BRET pairings with stimulation of PAR4 by the tethered ligand-mimicking peptide AYPGKF-NH_2_. Curve fitting was conducted by three-parameter, non-linear regression and EC_50_ values are reported for each G protein subtype. Where curve fitting could not be conducted EC_50_ values are reported as “not determined” (*n.d.*). Technical replicates were collected in triplicate for each experimental replicate (*n = 3-4*)

**Fig. 2.**
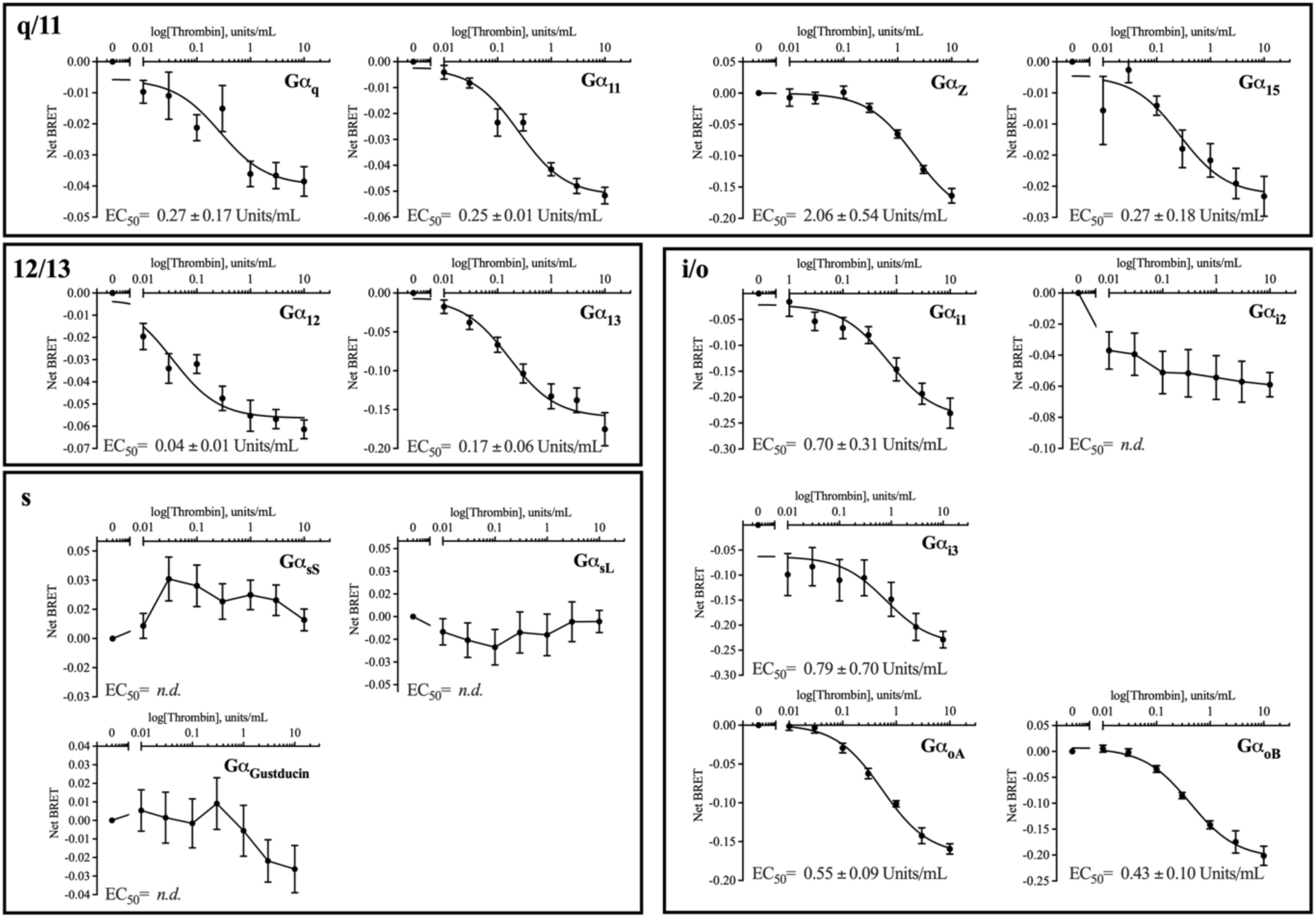
Thrombin-activated PAR4 stimulates Gα_q/11_, Gα_12/13_, and Gα_i/o_ G protein-subtypes, but not Gα_s_ family subtypes. Assays with PAR1-KO-HEK-293 cells expressing wild-type PAR-YFP receptor were conducted for each of the TRUPATH BRET pairings with stimulation of PAR4 by enzymatic activation with thrombin. Curve fitting was conducted by three-parameter, non-linear regression and EC_50_ values are reported for each G protein subtype. Where curve fitting could not be conducted EC_50_ values are reported as “not determined” (*n.d.*). Technical replicates were collected in triplicate for each experimental replicate (*n = 3-4*).

Overall, here we established the G protein subtypes that can be activated by PAR4 in HEK-293 and PAR1-knockout HEK-293 cells and our findings are consistent with well-established Gα_q/11_, Gα_12/13_, Gα_i_ signalling reported following PAR4 activation in physiologically relevant cell types^7,12,19,23^. With the complete PAR4 G protein coupling profile in hand, we next turned to examining the effect of mutations to PAR H8 and CT residues on G protein and β-arrestin recruitment.

### Membrane expression of wild-type and mutant PAR4-YFP receptors

In certain Class A GPCRs, H8 and CT mutations analogous to those discussed in the remainder of this study negatively impacted cell surface expression. For instance, Mutation of H8 lysine residues reduce membrane expression of both MCH_1_R and B_2_R ^24,25^. Additionally, mutation of the highly-conserved NPxxYx_5,6_F motif in rhodopsin and β_2_AR significantly affect cell surface expression of these receptors ^26–29^. Previously, we demonstrated that H8 R^352^AGLFQRS^359^ deletion did not impact appropriate cell membrane localization of PAR4 ^30^. Moreover, truncation studies wherein the CT of PAR4 is removed from H8 Lys^350^ (ΔK350) or CT Lys^367^ (ΔK350) reported appropriate cell membrane expression and agonist-dependent internalization ^31^. We therefore assessed cell surface expression of PAR4 H8 and CT mutants and observed no deficits in cell membrane expression compared to the wild-type receptor (Supplementary Figure 1).

### AYPGKF-NH_2_- and thrombin-stimulated G protein signalling is dependent on H8 R^352^AGLFQRS^359^ residues

Previously, we reported that the deletion of PAR4 H8 residues R^352^AGLFQRS^359^ (construct designated dRS-PAR4) exhibited defects in agonist-stimulated Gα_q/11_-mediated calcium signalling^30^. Here we employed a more conservative approach through alanine scanning mutations to rule out the potential for structural changes introduced by deletion contributing to the previously observed signalling defects.

PAR4 activation-triggered calcium signalling was measured as a proxy for Gα_q/11_ activation by PAR4 since we have previously demonstrated that the Gα_q/11_ inhibitor YM254890 completely abolished PAR4 calcium responses ^32^ and calcium signaling provided a more consistent means of assessing Gα_q/11_ mediated signaling compared to the TRUPATH G protein activation assay (Supplementary Figures 2 and 3). We find that mutation of the H8 RAGLFQRS motif to alanine (PAR4^R352-S359A^-YFP) significantly decreased both AYPGKF-NH_2_- (EC_50_ = *n.d.*, Max. = 2.5 ± 2.5% A23187, where “Max.” represents mean ± SEM of percent of A23187 calcium signal at 300 µM AYPGKF-NH_2_ or 10 units/mL thrombin; Fig. 2A) and thrombin-stimulated (EC_50_ = *n.d.*, Max. = 6.1 ± 1.0% A23187, Fig. 3H) calcium signalling compared to wild-type PAR4 (AYPGKF-NH_2_ EC_50_ = 26.7 ± 9.6 µM, Max. = 26.3 ± 2.5% A23187, Fig. 3A; thrombin EC_50_ = 0.3 ± 0.1 units/mL, Max. = 14.8 ± 1.4% A23187, Fig. 3H). Building on our previous findings, we investigated the impact of R^352^AGLFQRS^359^ to alanine mutation, on G protein activation by PAR4 (Fig. 1 and Fig. 2; Gα_12/13_ subtypes represented by Gα_13_ TRUPATH biosensor and Gα_i/o_ subtypes represented by Gα_oB_ TRUPATH biosensor). Interestingly, while substantial deficits were observed in Gα_q/11_-mediated calcium signalling, we did not see any significant alterations to PAR4^R352-S359A^-YFP mediated Gα_13_ or Gα_oB_ activation compared to wild-type receptor when stimulated with either thrombin or AYPGKF-NH_2_ (PAR4^R352-S359A^-YFP Gα_13_: AYPGKF-NH_2_ EC_50_ = 20.3 ± 6.4 µM, Max. = -0.18 ± 0.01 net BRET, Fig. 3B; Thrombin EC_50_ = 2.5 ± 0.8 units/mL, Max. = -0.17 ± 0.02 net BRET, Fig. 3I) (PAR4^R352-S359A^-YFP Gα_oB_: AYPGKF-NH_2_ EC_50_ = 25.3 ± 14.6 µM, Max. = - 0.12 ± 0.02 net BRET, Fig. 3C; Thrombin EC_50_ = *n.d.*, Max. = -0.39 ± 0.27 net BRET, Fig. 3J) (WT PAR4-YFP Gα_13_: AYPGKF-NH_2_ EC_50_ = 15.6 ± 2.5 µM, Max. = -0.17 ± 0.01 net BRET, Fig. 3B; Thrombin EC_50_ = 1.6 ± 0.3 units/mL, Max. = -0.14 ± 0.01 net BRET, Fig. 3I) (WT PAR4-YFP Gα_oB_: AYPGKF-NH_2_ EC_50_ = 27.4 ± 7.6 µM, Max. = -0.16 ± 0.01 Net BRET, Fig. 3C; Thrombin EC_50_ = 5.4 ± 3.3 units/mL, Max. = -0.17 ± 0.05 Net BRET, Fig. 3J).

**Fig. 3.**
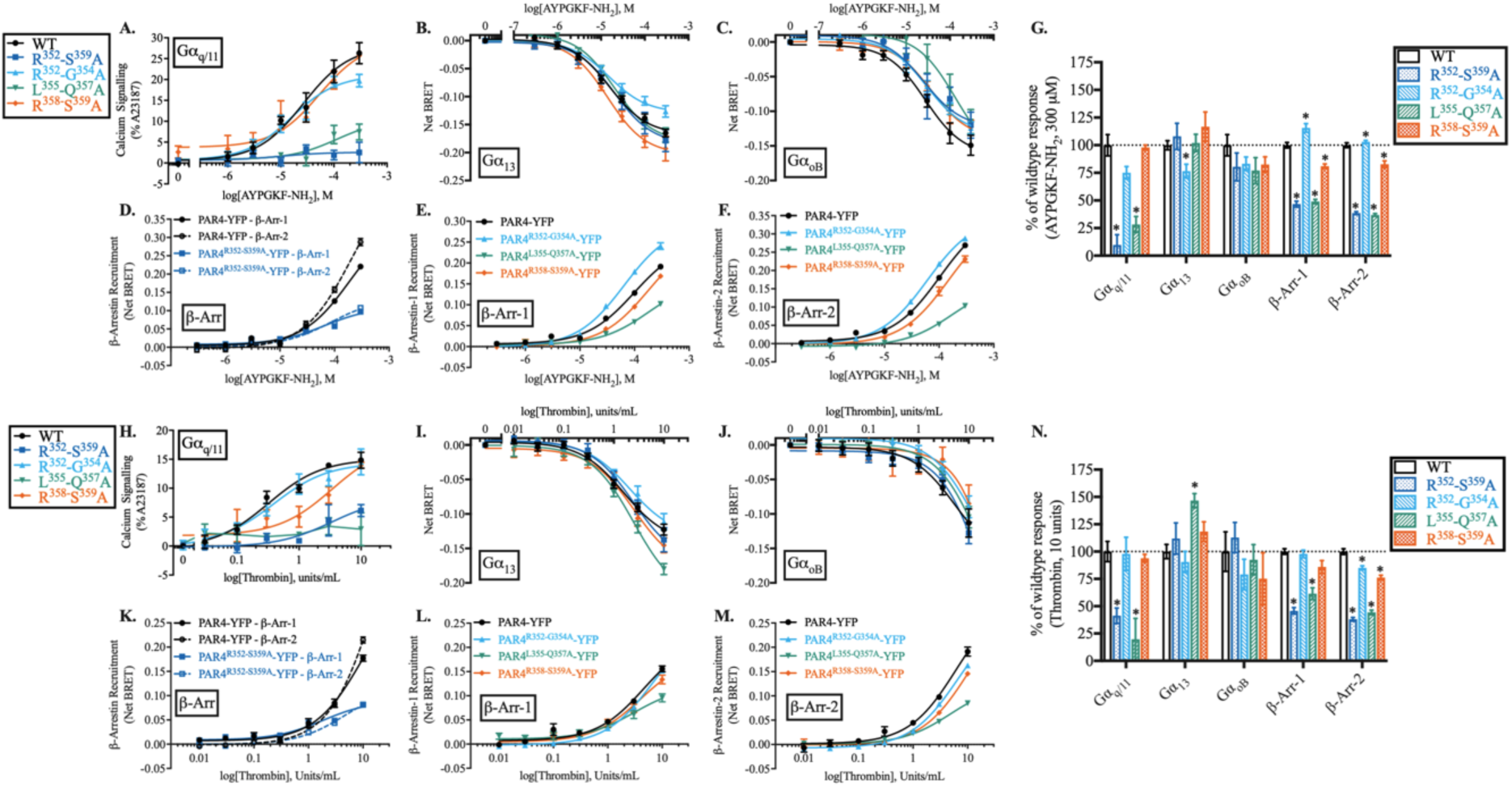
H8 Lys^355^-Glu^357^ residues are essential for Gα_q/11_ activation and β-arrestin-1/-2 recruitment to both AYPGKF-NH_2_- and thrombin-stimulated PAR4. *G protein activation (A-C, H-J).* Calcium signalling from wild-type PAR4-YFP and R^352^AGLFQRS^359^ total or sequential mutant PAR4 receptors in response to AYPGKF-NH_2_ (A) or thrombin (H) stimulation. Nonlinear regression curve fits are shown (mean ± S.E.) for three independent experiments (*n = 3*). G protein activation (TRUPATH) was recorded in cells expressing wild-type, R^352^AGLFQRS^359^ total or sequential mutant PAR4 receptors in response to either AYPGKF-NH_2_ [HEK-293 cells; Gα_13_ (B), Gα_oB_ (C)] or thrombin [PAR1-KO-HEK-293 cells; Gα_13_ (I), Gα_oB_ (J)]. Nonlinear regression curve fits are shown (mean ± S.E.). Technical replicates were collected in triplicate for each experimental replicate (*n = 3-4*). β*-arrestin recruitment (D-F, K-M).* Recruitment of β-arrestin-1/-2 to PAR4-YFP wild-type or R^352^AGLFQRS^359^Ala mutant PAR4 following stimulation with AYPGKF-NH_2_ (D) or thrombin (K). Recruitment of β-arrestin-1 (E) or -2 (F) to wild-type and sequential R^352^AGLFQRS^359^ mutant receptors in response to AYPGKF-NH_2_. Recruitment of β-arrestin-1 (L) or -2 (M) to wild-type and sequential R^352^AGLFQRS^359^ mutant receptors in response to thrombin stimulation. Nonlinear regression curve fits are shown (mean ± S.E.) for three to four independent experiments with triplicate data points for each concentration and receptor collected within each experiment. (*n = 3-4)*. *Summary data normalized to wild-type receptor (G, N)*. For ease of comparison, signalling of mutant PAR4 receptors was normalized to signalling recorded from wild-type PAR4 receptor in response to AYPGKF-NH_2_ (G) or thrombin (N) at the highest concentrations tested (300 µM AYPGKF-NH_2_, 10 units/mL thrombin) for each of the G protein activation and β-arrestin recruitment assays (A-F, H-M). Significance was calculated on raw (non-normalized) values for each assay by T-test and is indicated (*p < 0.05 compared to wild-type PAR4-YFP receptor).

To further delineate H8 residues involved in PAR4-mediated G protein signalling, we recorded agonist-stimulated Gα_q/11_-mediated calcium signalling and Gα_13_ and Gα_oB_ TRUPATH sensor BRET in cells expressing PAR4 receptor mutants with sequential H8 mutations. Mutation of Arg^352^-Gly^354^Ala did not significantly alter AYPGKF-NH_2_ or thrombin-stimulated calcium signalling (AYPGKF-NH_2_ EC_50_ = 15.6 ± 5.0 µM, Max. = 19.7 ± 1.5% A23187, Fig. 3A; thrombin EC_50_ = 0.4 ± 0.2 units/mL, Max. = 14.4 ± 1.2% A23187, Fig. 3H) compared to PAR4-YFP. Further, we observed no difference in Gα_oB_ signalling with either agonist (AYPGKF-NH_2_ EC_50_ = 29.6 ± 8.1 µM, Max. = -0.13 ± 0.01 net BRET, Fig. 3C; thrombin EC_50_ = 13.2 ± 13.0 units/mL, Max. = -0.22 ± 0.14 net BRET, Fig. 3J] and only observed a modest decrease in maximal signalling with AYPGKF-NH_2_-stimulated Gα_13_ activation [AYPGKF-NH_2_ EC_50_ = 15.1 ± 3.5 µM, Max. = - 0.13 ± 0.01 net BRET (p < 0.001), Fig. 3C; thrombin EC_50_ = 2.1 ± 0.6 units/mL, Max. = -0.13 ± 0.01 net BRET, Fig. 3J].

Similarly, there was no significant difference in Gα_q/11_-mediated calcium signalling with Arg^358^-Ser^359^Ala mutation in response to either AYPGKF-NH_2_ (EC_50_ = *n.d.*, Max. = 25.8 ± 0.6% A23187, Fig. 3A) or thrombin (EC_50_ = *n.d.*, Max. = 13.9 ± 0.5% A23187, Fig. 3H). We observed increased Gα_13_ maximal signalling from PAR4^R358-S359A^-YFP [AYPGKF-NH_2_ EC_50_ = 14.1 ± 3.1 µM, Max. = -0.20 ± 0.01 net BRET (p < 0.05), Fig. 3B; Thrombin EC_50_ = 2.5 ± 0.8 units/mL, Max. = -0.18 ± 0.02 net BRET (p = 0.04), Fig. 3I]; however, when comparing the net BRET values at the highest concentrations tested, there were no significant differences between the PAR4^R358-S359A^ mutant and wild-type receptor [Net BRET at AYPGKF-NH_2_ (300 µM) PAR4^R358-S359A^-YFP -0.19 ± 0.02 (p = 0.27), Fig. 3G; net BRET at thrombin (10 units/mL) PAR4^R358-S359A^-YFP -0.15 ± 0.01 (p = 0.11), Fig. 3N]. Similarly, we observed no significant impact of Arg^358^-Ser^359^Ala mutation on Gα_oB_ (AYPGKF-NH_2_ - EC_50_ = 37.4 ± 11.9 µM, Max. = -0.14 ± 0.01 net BRET, Fig. 3C; Thrombin EC_50_ = *n.d.*, Max. = *n.d.*, net BRET at 10 units/mL = -0.08 ± 0.03 (p = 0.42), Fig. 3J & 3N] signalling compared to wild-type receptor.

Interestingly, mutation of Leu^355^-Gln^357^ to alanine significantly decreased thrombin stimulated calcium signalling (EC_50_ = *n.d.*, Max. = 7.4 ± 1.9% A23187, Fig. 3A) (EC_50_ = *n.d.*, Max. = 2.9 ± 2.8% A23187, Fig. 3H) but, did not impact AYPGKF-NH_2_ Gα_13_ (EC_50_ = 20.2 ± 4.1 µM, Max. = -0.18 ± 0.01 net BRET, Fig. 3B) or Gα_oB_ (EC_50_ = *n.d.*, Max. = *n.d.*, net BRET at 300 µM = -0.11 ± 0.02, Fig. 3C) activation. We observed that the Leu^355^-Gln^357^Ala mutation did modestly increase thrombin-stimulated Gα_13_ maximal net BRET [EC_50_ = 2.5 ± 0.5 units/mL, Max. = -0.22 ± 0.02 net BRET (p < 0.001), Fig. 3I] but had no impact on Gα_oB_ activation (EC_50_ = 17.0 ± 16.0 units/mL, Max. = -0.28 ± 0.18 net BRET, Fig. 3J).

Overall, we conclude that the loss of calcium signalling, previously reported with the H8 R^352^AGLFQRS^359^ motif deletion^30^ or alanine substitution in the current study, is not due to truncation or shortening of the CT, rather, the deficits in calcium signalling appear to be mediated through loss of effector interactions with this site. The H8 Leu^355^-Gln^357^ residues specifically are involved in agonist-stimulated Gα_q/11_ coupled calcium signalling downstream of activated PAR4. Further, these residues are important for both synthetic peptide and thrombin revealed tethered ligand mediated signaling from PAR4. The modest impact of Leu^355^-Gln^357^Ala mutation on Gα_13_ and Gα_oB_ also suggest that these residues may be involved in a G protein subtype specific interaction of Gα_q/11_ with PAR4.

### AYPGKF-NH_2_ and thrombin stimulated PAR4 β-arrestin-1 and -2 recruitment involves H8 Leu^355^-Gln^357^

Previously, we reported that in addition to defects in agonist-stimulated Gα_q/11_ coupled calcium signalling, deletion of the H8 R^352^AGLFQRS^359^ motif (dRS-PAR4) decreased β-arrestin recruitment to PAR4^30^. As with calcium signalling, we sought to determine the role of H8 mutations on β-arrestin-1/-2 recruitment in response to either peptide- or enzyme-mediated activation of PAR4. As previously demonstrated, the concentration effect curve for β-arrestin-1/-2 recruitment to PAR4 does not saturate in response to either peptide or thrombin activation, therefore the data presented throughout for β-arrestin recruitment are the net BRET ratio at the highest concentration tested for each agonist for comparison between receptor mutants (AYPGKF-NH_2_ 300 µM, thrombin 10 units/mL). In response to AYPGKF-NH_2_-stimulation, we observed β-arrestin-1 (Max. = 0.22 ± 0.01 net BRET; where “Max.” represents mean ± SEM of net BRET at 300 µM AYPGKF-NH_2_) and β-arrestin-2 (Max. = 0.29 ± 0.01 net BRET) recruitment consistent with previous findings (Fig. 3D-F) ^13,30^. Thrombin-stimulated recruitment of β-arrestin-1 (Max. = 0.18 ± 0.01 net BRET; where “Max.” represents mean ± SEM of net BRET at 10 units/mL thrombin) and β-arrestin-2 (Max. = 0.21 ± 0.01 net BRET) was also recorded (Fig. 3K-M) and as previously observed, recruitment of β-arrestin-1/-2 in response to thrombin exhibits decreased maximal recruitment compared to peptide activation of the receptor^17,30^.

Mutation of the entire H8 R^352^AGLFQRS^35^ sequence to alanine (PAR4^R352-S359A^-YFP) mirrored the findings of our previous results using a truncation mutant providing confidence that the residues, and not the deletion, were responsible for the effects previously observed. β-arrestin- 1/-2 recruitment to PAR4^R352-S359A^-YFP was significantly decreased in response to both AYPGKF-NH_2_ [β-Arr-1 Max. 0.10 ± 0.01 net BRET (p < 0.0001); β-Arr-2 Max. 0.11 ± 0.00 net BRET (p < 0.0001), Fig. 3D] and thrombin [β-Arr-1 Max. 0.08 ± 0.01 net BRET (p < 0.0001); β-Arr-2 Max. 0.08 ± 0.00 net BRET (p < 0.0001), Fig. 3K] stimulation indicating that these H8 residues are indeed important for β-arrestin-1/-2 recruitment to PAR4.

To determine whether the entire H8 R^352^AGLFQRS^359^ sequence, or a subset of these residues, is participating in agonist-stimulated recruitment of β-arrestins, we utilized the sequential H8 mutant receptors and recorded recruitment. The first of these mutations, Arg^352^-Gly^354^Ala, significantly increased recruitment of β-arrestin-1 [Max. = 0.24 ± 0.01 net BRET (p < 0.05)] and -2 [Max. = 0.29 ± 0.00 (p < 0.05)] compared to wild-type PAR4-YFP in response to AYPGKF-NH_2_ stimulation (Fig. 3E & 3F). Interestingly, we observed differential agonist-dependent recruitment of β-arrestins to thrombin activated PAR4^R352-G354A^-YFP. Thrombin-stimulated recruitment of β-arrestin-1 to PAR4^R352-G354A^-YFP [Max. = 0.15 ± 0.01 net BRET (p = 0.69), Fig. 3L] was not significantly different than wild-type receptor, however, the recruitment of β-arrestin-2 was significantly reduced [Max. = 0.16 ± 0.00 net BRET (p = 0.01), Fig. 3M]. Therefore, these data may highlight differences in mechanisms underlying β-arrestin-1and 2 recruitment to PAR4.

Leu^355^-Gln^357^ to alanine mutation (PAR4^L355-Q357A^-YFP) resulted in decreased β-arrestin-1/-2 recruitment to PAR4 in response to both agonists studied. AYPGKF-NH_2_-stimulated recruitment of β-arrestin-1 [Max. = 0.10 ± 0.00 net BRET (p < 0.0001), Fig. 3E) and -2 [Max. = 0.10 ± 0.00 net BRET (p < 0.0001), Fig. 3F) to PAR4^L355-Q357A^-YFP was significantly reduced compared to wild-type receptor. Further, thrombin-stimulated recruitment of β-arrestin-1 [Max. = 0.10 ± 0.01 net BRET (p < 0.0001), Fig. 3L) and -2 [Max. = 0.08 ± 0.00 net BRET (p < 0.0001), Fig. 3M) was also significantly reduced compared to thrombin-activated PAR4. Given that both peptide- and thrombin-stimulated recruitment of β-arrestins was affected by Leu^355^-Gln^357^Ala mutation, we determine a role for these residues in β-arrestin recruitment to PAR4, independent of agonist.

Finally, we evaluated the effect of alanine mutation Arg^358^ and Ser^359^ (PAR4^R358-S359A^-YFP). We observed that AYPGKF-NH_2_-stimulated recruitment of both β-arrestin-1 [Max. = 0.17 ± 0.00 net BRET (p < 0.05), Fig. 3E] and -2 [Max. = 0.23 ± 0.01 net BRET (p < 0.05), Fig. 3F) was significantly reduced compared to wild-type receptor. Interestingly, as observed with Arg^352^-Gly^354^Ala mutation, thrombin-stimulated β-arrestin-1 recruitment was unaffected with Arg^358^-Ser^359^Ala mutation [Max. = 0.13 ± 0.01 net BRET (p = 0.06), Fig. 3L); however, β-arrestin-2 recruitment was significantly reduced [Max. = 0.15 ± 0.00 net BRET (p < 0.05), Fig. 3M).

These data highlight an important role for H8 R^352^AGLFQRS^359^ residues in the recruitment of β-arrestins to PAR4. Sequential mutation revealed a role for Leu^355^-Gln^357^ residues in agonist-stimulated β-arrestin recruitment, regardless of which PAR4-agonist was applied. Thus, these residues may be a key regulatory site for this interaction as the phenotype is observed in response to either agonist tested. Interestingly, the data also reveal key agonist-dependent differences in which residues are important for β-arrestin recruitment. Peptide-stimulated β-arrestin-1/-2 recruitment to PAR4 was altered by all of the H8 mutations studied, with Leu^355^-Gln^357^Ala mutation more deleterious than the other mutations studied (p < 0.05). Interestingly, thrombin-stimulated recruitment of β-arrestin-1/-2 was also significantly decreased compared to wild-type receptor, like the effect observed with peptide stimulation of this mutant. All mutations studied were significantly deleterious to β-arrestin-2 recruitment downstream of thrombin-activation of the receptor; however, only Leu^355^-Gln^357^Ala mutation altered thrombin-stimulated recruitment of β-arrestin-1 in comparison to wild-type receptor. Thus, these results indicate both an agonist-dependent, and β-arrestin-subtype dependent role for H8 residues in recruitment to activated PAR4.

### AYPGKF-NH_2_ or thrombin stimulated PAR4 activation of Gα_q/11_ and Gα_oB_ signalling requires H8 Lys^350^

H8 and C-terminal tail lysine residues have been shown for many Class A GPCRs to be involved in interactions with both G proteins and β-arrestins, as well as changes in the activation state of GPCRs ^33–35^. Specifically, lysine residues located in the H8, which are adjacent to the canonical NPxxYX_5,6_F motif, have been implicated to be a binding site for various G protein subtypes including Gα_t_, Gα_i_, and Gα_s_ ^24,25,35,36^. The PAR4 H8 and C-terminal region contains two lysine residues, Lys^350^ (8.53, Ballesteros-Weinstein numbering) and Lys^367^, which are distal to the PAR4 DPxxYX_5,6_F motif. Given the emerging role for H8 lysine residues in GPCR-G protein interaction, we investigated G protein activation in PAR4 receptors with C-terminal tail lysine mutations. Gα_q/11_-mediated calcium signalling in HEK-293 cells expressing PAR4-YFP was recorded in response to AYPGKF-NH_2_- (EC_50_ = 35.9 ± 12.8 µM, Max. = 26.1 ± 2.6% A23187, Fig. 4A) or thrombin-stimulation (EC_50_ = 1.3 ± 0.5 units/mL, Max. = 16.2 ± 1.7% A23187, Fig. 4H). In comparison, cells expressing PAR4 with H8 lysine, Lys^350^Ala mutation (PAR4^K350A^-YFP) had significantly decreased calcium signalling in response to both AYPGKF-NH_2_ (EC_50_ = *n.d.*, Max. at 300 µM = 8.5 ± 1.8% A23187; Fig. 4A) and thrombin (EC_50_ = *n.d.*, Signal at 300 µM = 4.3 ± 0.7% A23187, Fig. 4H).

**Fig. 4.**
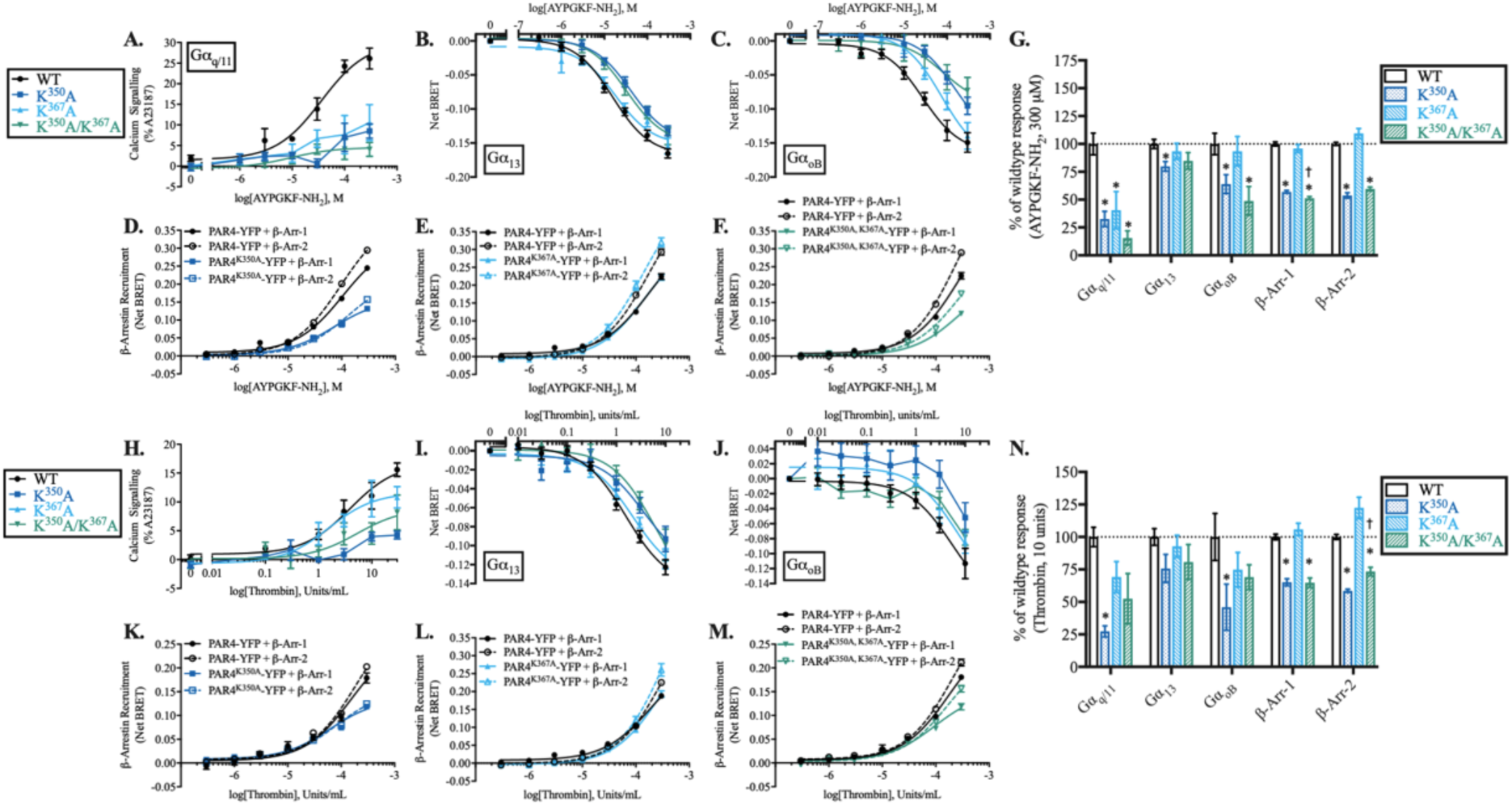
H8 Lys^350^ residue is essential for Gα_q/11_ and Gα_oB_ activation and β-arrestin-1/-2 recruitment to both AYPGKF-NH_2_- and thrombin-stimulated PAR4, but not Gα_13_ activation. *G protein activation (A-C, H-J).* Calcium signalling from wild-type PAR4-YFP, PAR4^K350A^-YFP, PAR4^K367A^-YFP, and PAR4^K350A, K367A^-YFP receptors in response to AYPGKF-NH_2_ (A) or thrombin (H) stimulation. Nonlinear regression curve fits are shown (mean ± S.E.) for three independent experiments (*n = 3-4*). G protein activation (TRUPATH) was recorded in cells expressing wild-type or mutant PAR4-YFP receptors in response to either AYPGKF-NH_2_ [HEK- 293 cells; Gα_13_ (B), Gα_oB_ (C)] or thrombin [PAR1-KO-HEK-293 cells; Gα_13_ (I), Gα_oB_ (J)]. Nonlinear regression curve fits are shown (mean ± S.E.). Technical replicates were collected in triplicate for each experimental replicate (*n = 3-4*). β*-arrestin recruitment (D-F, K-M).* Recruitment of β-arrestin-1/-2 to PAR4-YFP wild-type or Lys^350^Ala mutant PAR4-YFP following stimulation with AYPGKF-NH_2_ (D) or thrombin (K). Recruitment of β-arrestin-1/-2 to PAR4- YFP wild-type or Lys^367^Ala mutant PAR4-YFP following stimulation with AYPGKF-NH_2_ (E) or thrombin (L). Recruitment of β-arrestin-1/-2 to PAR4-YFP wild-type or Lys^350^Ala/Lys^367^Ala double mutant PAR4-YFP following stimulation with AYPGKF-NH_2_ (F) or thrombin (M). Nonlinear regression curve fits are shown (mean ± S.E.) with triplicate data points for each concentration and receptor collected within each experiment. (*n = 3)*. *Summary data normalized to wild-type receptor (G, N)*. For ease of comparison, signalling of mutant PAR4 receptors was normalized to signalling recorded from wild-type PAR4 receptor in response to AYPGKF-NH_2_ (G) or thrombin (N) at the highest concentrations tested (300 µM AYPGKF-NH_2_, 10 units/mL thrombin) for each of the G protein activation and β-arrestin recruitment assays (A-F, H-M). Significance was calculated on raw (non-normalized) values for each assay by T-test and is indicated (*p < 0.05 compared to wild-type, ^†^p < 0.05 compared to PAR4^K367A^-YFP).

Additionally, we observed that Lys^350^Ala mutation decreased both AYPGKF-NH_2_-stimulated Gα_13_ [EC_50_ = 41.3 ± 9.2 µM (p < 0.05), Max. = -0.15 ± 0.01 net BRET (p = 0.12), net BRET at 300 µM = -0.13 ± 0.01 (p < 0.05), Fig. 4B] and Gα_oB_ [EC_50_ = *n.d.*, Max. = *n.d.*, net BRET at 300 µM = -0.10 ± 0.01 (p < 0.05), Fig. 4C] activation compared to wild-type receptor [Gα_13_ - EC_50_ = 15.6 ± 1.5 µM, Max. = -0.17 ± 0.01 net BRET, Fig. 4B; Gα_oB_ - EC_50_ = 27.4 ± 7.6 µM, Max. = -0.16 ± 0.01 net BRET, Fig. 4C]. In contrast, Lys^350^Ala mutation significantly impacted thrombin-stimulated Gα_oB_ signalling [EC_50_ = *n.d.*, Max. = *n.d.*, net BRET at 10 units/mL = -0.05 ± 0.02 (p < 0.05), Fig. 4J] but not Gα_13_ signalling [EC_50_ = 3.6 ± 2.1 units/mL (p = 0.13), Max. = - 0.12 ± 0.03 net BRET (p = 0.50), net BRET at 10 units/mL = -0.09 ± 0.01 (p = 0.07), Fig. 4I] compared to wild-type receptor [Gα_13_ - EC_50_ = 1.6 ± 0.3 units/mL, Max. = -0.14 ± 0.01 net BRET, Fig. 4I; Gα_oB_ - EC_50_ = 5.4 ± 3.3 units/mL, Max. = -0.17 ± 0.05 net BRET, Fig. 4J].

Mutation of the distal C-terminal lysine residue (PAR4^K367A^-YFP) also decreased calcium signalling following receptor stimulation with either AYPGKF-NH_2_ [EC_50_ = *n.d.*, Signal at 300 µM = 10.6 ± 4.4% A23187 (p < 0.05); Fig. 4.A] or thrombin [EC_50_ = 0.5 ± 0.3 units/mL, Max. = 10.8 ± 1.9% A23187 (p < 0.05); Fig. 4H]. In contrast to the decreased signalling observed with Lys^350^Ala mutation, Lys^367^Ala mutation did not significantly alter either AYPGKF-NH_2_- or thrombin-stimulated Gα_13_ [AYPGKF-NH_2_ - EC_50_ = 18.3 ± 5.4 µM (p = 0.67), Max. = -0.15 ± 0.01 net BRET (p = 0.25), Fig. 4B; Thrombin - EC_50_ = 2.2 ± 0.6 units/mL (p = 0.29), Max. = -0.14 ± 0.01 net BRET (p = 0.66), Fig. 4I] or Gα_oB_ [AYPGKF-NH_2_ - EC_50_ = 75.7 ± 26.4 µM (p < 0.05), Max. = -0.17 ± 0.02 net BRET (p = 0.65), Fig. 4C; Thrombin - EC_50_ = 6.8 ± 6.5 units/mL (p = 0.82), Max. = -0.16 ± 0.08 net BRET (p = 0.87), Fig. 4J] activation.

Combination of these lysine mutations (PAR4^K350A, K367A^-YFP) also significantly reduced AYPGKF-NH_2_-stimulated calcium signalling [EC_50_ = *n.d.*, Max. = *n.d.*, Signal at 300 µM = 4.0 ± 1.7% A23187 (p < 0.05), Fig. 4A) but not thrombin-stimulated signalling compared to wild-type [EC_50_ = 1.8 ± 2.0 units/mL (p = 0.79), Max. = 9.1 ± 2.8% A23187 (p = 0.22), Signal at 300 µM = 8.2 ± 3.0 (p = 0.08), Fig. 4H]. We observed that Gα_13_ was able to reach equivalent signalling by the highest concentrations of agonist tested however revealed a distinct rightward shift in the concentration of agonist required to reach EC_50_ with both AYPGKF-NH_2_ [EC_50_ = 32.8 ± 8.8 µM (p < 0.05), Max. = -0.15 ± 0.01 net BRET (p = 0.20), Fig. 4C] and thrombin [EC_50_ = 6.0 ± 3.7 µM (p < 0.05), Max. = -0.16 ± 0.04 net BRET (p = 0.63), Fig. 4I]. In contrast, Gα_oB_ activation was significantly diminished compared to wild-type receptor with AYPGKF-NH_2_ stimulation [EC_50_ = *n.d.*, Max. = *n.d.*, net BRET at 300 µM = -0.07 ± 0.02 (p < 0.05), Fig. 4C] but not thrombin activation of PAR4 [EC_50_ = *n.d.*, Max. = *n.d.*, BRET at 10 units/mL = -0.08 ± 0.01 (p = 0.14), Fig. 4J].

Together, these data implicate a previously unreported role of H8 Lys^350^ in PAR4 Gα_q/11_-, Gα_13_-, and Gα_oB_-mediated signalling downstream of PAR4 activation with either agonist peptide-mediated or thrombin cleavage-mediated activation of calcium signalling. Whether the residue is involved directly in G protein effector binding or a structural interaction with other active state receptor motifs, such as the NPxxYx_4,5_F motif or through receptor post-translational modifications (e.g. ubiquitination), remains to be explored. Additionally, the role of the more distal Lys^367^ in peptide-activated PAR4 may provide some additional insight into key differences in effector signalling between peptide-activated and thrombin-activated PAR4 given that the effect of mutation was significantly deleterious to peptide agonist-stimulated, but not thrombin-stimulated, activation of Gα_q/11_ and Gα_oB_ (Fig. 4G, 4N). Further, these data may reveal G protein subtype-specific roles for Lys^367^ as Gα_13_ was unaffected with Lys^367^Ala mutation in response to either agonist.

### Importance of Lys^350^ in AYPGKF-NH_2_- and thrombin-stimulated β-arrestin-1 and -2 recruitment revealed by PAR4 CT lysine mutants

Lysine residues located in the H8, adjacent to the canonical NPxxYX_5,6_F motif, have also been implicated to have a role in interactions with the β-arrestin finger-loop, assisting the β-arrestin proteins in recognizing an active conformation of the receptor. To determine if these C-terminal tail lysine residues play a role in β-arrestin recruitment to activated PAR4, we recorded β-arrestin recruitment to PAR4^K350A^-YFP and PAR4^K367A-YFP^. Lys^350^Ala significantly reduced agonist-stimulated β-arrestin recruitment to PAR4 in response to AYPGKF-NH_2_ stimulation [PAR4^K350A^-YFP, β-Arr-1 Max. = 0.13 ± 0.00 net BRET (p < 0.0001), β-Arr-2 Max. = 0.16 ± 0.01 net BRET (p < 0.0001)] compared to wild-type PAR4-YFP (β-Arr-1 Max. = 0.24 ± 0.00 net BRET, β-Arr-2 Max. = 0.29 ± 0.01 net BRET, Fig. 4D). Additionally, thrombin-stimulated β-arrestin recruitment was also significantly decreased to PAR4^K350A^-YFP [β-Arr-1 Max. = 0.12 ± 0.00 net BRET (p < 0.0001), β-Arr-2 Max. = 0.12 ± 0.00 net BRET (p < 0.0001)] compared to wild-type receptor (β-Arr-1 Max. = 0.18 ± 0.01 net BRET, β-Arr-2 Max. = 0.20 ± 0.00 net BRET, Fig. 4K). As discussed above, the CT of PAR4 contains one additional, non-H8 lysine residue, Lys^367^. Lys^367^Ala mutation had no effect on agonist-stimulated β-arrestin recruitment to the receptor compared to wild-type receptor in response to either AYPGKF-NH_2_ [β-Arr-1 Max. = 0.22 ± 0.01 net BRET (p = 0.85), β-Arr-2 Max. = 0.32 ± 0.01 net BRET (p = 0.10), Fig. 4E] or thrombin [β-Arr-1 Max. = 0.19 ± 0.01 net BRET (p = 0.60), β-Arr-2 Max. = 0.26 ± 0.02 net BRET (0.07), Fig. 4L] compared to wild-type receptor (AYPGKF-NH_2_ β-Arr-1 Max. = 0.22 ± 0.01 net BRET, β-Arr-2 Max. = 0.29 ± 0.01 net BRET, Fig. 4E; thrombin β-Arr-1 Max. = 0.19 ± 0.01 net BRET, β-Arr-2 Max. = 0.23 ± 0.01 net BRET, Fig. 4L).

As expected given the effects of Lys^350^Ala mutation on β-arrestin recruitment, combined mutation of both lysine residues (PAR4^K350A, K367A^-YFP) significantly decreased in response to AYPGKF-NH_2_ [β-Arr-1 Max. = 0.12 ± 0.00 net BRET (p < 0.0001), β-Arr-2 Max. = 0.17 ± 0.01 net BRET (p < 0.0001), Fig. 4F] or thrombin [β-Arr-1 Max. = 0.12 ± 0.01 net BRET (p < 0.0001), β-Arr-2 Max. = 0.16 ± 0.01 net BRET (p < 0.0001), Fig. 4M].

Interestingly, in response to peptide agonism of the double mutant receptor, β-arrestin-1 recruitment was significantly decreased when compared to Lys^350^Ala mutation alone (p < 0.05); however, β-arrestin-2 recruitment was not significantly different than single mutation of the H8 Lys^350^ (p = 0.07). Oppositely, thrombin-stimulated β-arrestin-1 recruitment did not show a significantly greater reduction than with the single Lys^350^Ala mutation alone (p = 0.95), however, β-arrestin-2 recruitment was statistically recovered compared to Lys^350^Ala mutation alone (p < 0.0001). Thus, the H8 Lys^350^ residue appears to have a role in agonist-stimulated β-arrestin recruitment. Additionally, there may be a role for the distal Lys^367^ in agonist-dependent or subtype specific differential recruitment of β-arrestins.

### DPxxYx_6_F motif residues, Tyr^340^ and Phe^347^, are necessary for appropriate activation of Gα_q/11_, Gα_13_, and Gα_oB_ following AYPGKF-NH_2_ or thrombin stimulation of PAR4

H8 and C-terminal tail residues are involved in regulating both interactions with β-arrestins and G proteins and changes in the activation state of many Class A GPCRs ^33–35^. Many Class A GPCRs possess the canonical NPxxYX_5,6_F motif spanning TM7 and H8, including residues Asn^7.49^, Pro^7.50^, Tyr^7.53^, and H8 Phe^8.50^. This motif is thought to be involved in stabilizing the inactive state of Class A GPCRs. Additionally, mutational studies have demonstrated deficits in G protein activation and signalling when Tyr^7.53^ and Phe^8.50^ are mutated to alanine ^26,37^. In PAR4, the NPxxYx_5,6_F motif is D^7.49^PFIY^7.53^YYVSAEF^8.50^. To determine if there is similar role for Tyr^7.53^ and Phe^8.50^ in PAR4-mediated activation of G proteins, single amino acid mutations of Tyr^340^ (PAR4^Y340A^-YFP) and Phe^347^ (PAR4^F347A^-YFP) to alanine were generated.

Gα_q/11_-mediated calcium signalling in HEK-293 cells expressing PAR4-YFP was recorded in response to AYPGKF-NH_2_- (EC_50_ = 19.4 ± 8.4 µM, Max. = 20.6 ± 1.8% A23187; Fig. 5A) or thrombin-stimulation (EC_50_ = 0.4 ± 0.2 units/mL, Max. = 19.4 ± 2.6% A23187; Fig. 5G). Cells expressing PAR4 with mutation of TM7, Tyr^340^Ala (PAR4^Y340A^-YFP) had significantly decreased calcium signalling in response to both AYPGKF-NH_2_ (EC_50_ = *n.d.*, Max. = *n.d.*, Signal at 300 µM= 0.9 ± 2.4% A23187 (p < 0.05), Fig. 5A] and thrombin [EC_50_ = *n.d.*, Max. = *n.d.*, Signal at 10 units/mL = 2.8 ± 0.5% A23187; Fig. 5G).

**Fig. 5.**
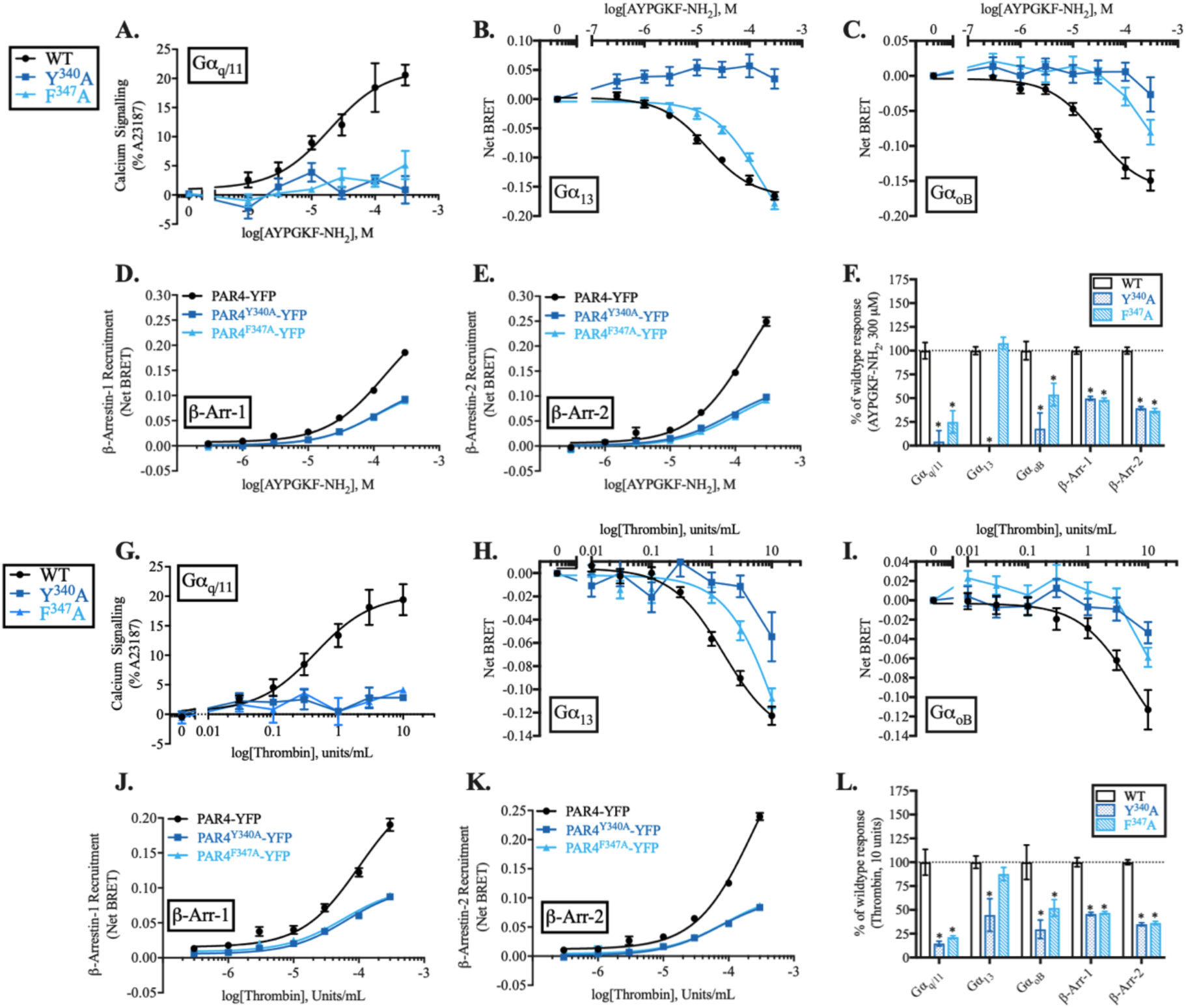
TM7 and H8 DPxxYx_6_F motif residues Tyr^340^ and Phe^347^ are essential for G protein activation and β-arrestin-1/-2 recruitment to both AYPGKF-NH_2_- and thrombin-stimulated PAR4, but not Phe^347^ in Gα_13_ activation. *G protein activation (A-C, H-J).* Calcium signalling from wild-type PAR4-YFP, PAR4^Y340A^-YFP, and PAR4^F347A^-YFP receptors in response to AYPGKF-NH_2_ (A) or thrombin (G) stimulation. Nonlinear regression curve fits are shown (mean ± S.E.) for three independent experiments (*n = 3*). G protein activation (TRUPATH) was recorded in cells expressing wild-type or mutant PAR4-YFP receptors in response to either AYPGKF-NH_2_ [HEK-293 cells; Gα_13_ (B), Gα_oB_ (C)] or thrombin [PAR1-KO-HEK-293 cells; Gα_13_ (H), Gα_oB_ (I)]. Nonlinear regression curve fits are shown (mean ± S.E.). Technical replicates were collected in triplicate for each experimental replicate (*n = 3-4*). β*-arrestin recruitment (D-E, J-K).* Recruitment of β-arrestins to wild-type or mutant PAR4-YFP receptors following stimulation with AYPGKF-NH_2_ [β-Arr-1 (D), β-Arr-2 (E)] or thrombin [β-Arr-1 (J), β-Arr-2 (K)]. Nonlinear regression curve fits are shown (mean ± S.E.) with triplicate data points for each concentration and receptor collected within each experiment. (*n = 3)*. *Summary data normalized to wild-type receptor (F, L)*. For ease of comparison, signalling of mutant PAR4 receptors was normalized to signalling recorded from wild-type PAR4 receptor in response to AYPGKF-NH_2_ (F) or thrombin (L) at the highest concentrations tested (300 µM AYPGKF-NH_2_, 10 units/mL thrombin) for each of the G protein activation and β-arrestin recruitment assays (A-E, G K). Significance was calculated on raw (non-normalized) values for each assay by T-test and is indicated (*p < 0.05 compared to wild-type PAR4-YFP receptor).

To determine whether the loss of function observed with PAR4^Y340A^-YFP is due to the loss of interaction with the H8 Phe^347^ residue, as would be expected as part of a traditional NPxxY_6_F motif, we generated a concomitant Phe^347^Ala mutant receptor and recorded calcium signalling. Like the deficits observed with Tyr^340^Ala mutation, PAR4^F347A^-YFP mutation significantly abrogated calcium signalling downstream of PAR4 activation with both AYPGKF-NH_2_ (EC_50_ = *n.d.*, Signal at 300 µM = 5.1 ± 2.4% A23187; Fig. 5A) and thrombin (EC_50_ = *n.d.*, Max. = 4.1 ± 0.3% A23187; Fig. 5G) stimulation. Thus, TM7 Tyr^340^ appears to perform an integral role in the DPxxYx_6_F motif in PAR4 Gα_q/11_-mediated calcium signalling through association with H8 Phe^347^.

To determine if the loss of Tyr^340^ or Phe^347^ impacted activation of other G proteins, as would be expected with a more global mechanism of PAR4 activation (as observed with many other Class A GPCRs), we recorded Gα_13_ and Gα_oB_ activation in response to either AYPGKF-NH_2_ or thrombin. Mutation of Tyr^340^ significantly abrogated both Gα_13_ and Gα_oB_ activation in response to either AYPGKF-NH_2_ [Gα_13_ - EC_50_ = *n.d.*, Max. = *n.d.*, net BRET at 300 µM = 0.03 ± 0.02 (p < 0.0001), Fig. 5B; Gα_oB_ - EC_50_ = *n.d.*, Max. = *n.d.*, net BRET at 300 µM = -0.03 ± 0.2 (p < 0.05), Fig. 5C] or thrombin [Gα_13_ - EC_50_ = *n.d.*, Max. = *n.d.*, net BRET at 10 units/mL = -0.05 ± 0.02 (p < 0.05), Fig. 5H; Gα_oB_ - EC_50_ = *n.d.*, Max. = *n.d.*, net BRET at 10 units/mL = -0.03 ± 0.1 (p < 0.05), Fig. 5I]. Interestingly, curve fitting of Gα_13_ activation in response to agonist was not possible for PAR4^F347A^-YFP mutation however, the net BRET achieved at the highest concentrations tested were equivalent to those achieved with wild-type receptor [AYPGKF-NH_2_ - EC_50_ = *n.d.*, Max. = *n.d.*, net BRET at 300 µM = -0.18 ± 0.01 (p = 0.32), Fig. 5B; Thrombin - EC_50_ = *n.d.*, Max. = *n.d.*, net BRET at 10 units/mL = -0.11 ± 0.01 (p = 0.21), Fig. 5H]. In contrast to Gα_13_, Gα_oB_ activation was significantly diminished in response to both agonists [AYPGKF-NH_2_ - EC_50_ = *n.d.*, Max. = *n.d.*, net BRET at 300 µM = -0.08 ± 0.02 (p < 0.05), Fig. 5C; Thrombin - EC_50_ = *n.d.*, Max. = *n.d.*, net BRET at 10 units/mL = -0.06 ± 0.01 (p < 0.05), Fig. 5I].

Together, these data demonstrate a role for the DPxxYx_6_F motif in PAR4 activation of G protein signalling. Notably, both mutations impacted G protein activation; however, both Gα_q/11_ and Gα_oB_ activation was all but completely abolished compared to wild-type receptor. In the present study we did not determine whether the decreased signalling is due to a lack of receptor-effector interaction or whether the receptor is in an active state upon loss of the Tyr^340^/Phe^347^ contact. We hypothesize that the latter is a likely mechanism given the evidence with many other Class A GPCRs in this contact mediating an inactive state of the receptor; thus, loss of this inactivating contact may place the receptor in a perpetually active state in which case further agonist-induced change in signalling are not seen. Further studies using PAR4 antagonists could determine if these mutations lead to constitutive activation of PAR4.

### The role of DPxxYx_6_F motif residues, Tyr^340^ and Phe^347^, in β-arrestin-1/-2 recruitment to activated PAR4

To determine whether the PAR4 NPxxYx_6_F motif contributes to β-arrestin-1/-2 recruitment we monitored β-arrestin recruitment to PAR4 receptors with Tyr^7.53^ (Tyr^340^Ala) and Phe^8.50^ (Phe^347^Ala) mutations. Tyr^340^Ala mutation significantly reduced agonist-stimulated β-arrestin recruitment to PAR4 in response to AYPGKF-NH_2_ stimulation [β-Arr-1 Max. = 0.09 ± 0.00 net BRET (p < 0.0001), β-Arr-2 Max. = 0.10 ± 0.00 net BRET (p < 0.0001)] compared to wild-type PAR4-YFP β-Arr-1 Max. = 0.19 ± 0.01 net BRET, β-Arr-2 Max. = 0.25 ± 0.01 net BRET, Fig. 5D & E). Interestingly, Phe^347^Ala mutation yielded almost superimposable deficits in peptide-stimulated β-arrestin recruitment, when compared to Tyr^340^Ala mutation [Phe^347^Ala, β-Arr-1 Max. = 0.09 ± 0.00 (p < 0.0001), β-Arr-2 Max. = 0.09 ± 0.01 (p < 0.0001)] (Fig. 5D & 5E).

Similar to the decreases observed with peptide stimulation, β-arrestin-1/-2 recruitment was significantly reduced in both PAR4^Y340A^-YFP [β-Arr-1 Max. = 0.09 ± 0.00 net BRET (p < 0.0001), β-Arr-2 Max. = 0.09 ± 0.00 net BRET (p < 0.0001)] and PAR4^F347A^-YFP (β-Arr-1 Max. = 0.08 ± 0.00net BRET (p < 0.0001), β-Arr-2 Max. = 0.09 ± 0.00 net BRET (p < 0.0001)] mutants compared to wild-type receptor (β-Arr-1 Max. = 0.19 ± 0.01 net BRET, β-Arr-2 Max. = 0.24 ± 0.01 net BRET) when stimulated with thrombin (Fig. 5J & 5K).

Given the significant reduction in β-arrestin recruitment observed with both thrombin- and peptide-stimulated PAR4 containing these mutations, Tyr^340^ and Phe^347^ are likely key residues involved in the TM7/H8 interactions governing β-arrestin recruitment. Indeed, interaction between these two residues as a part of the DPxxYx_6_F motif in PAR4 appears to be important for both G protein activation and β-arrestin recruitment.

### Impact of CT phosphorylatable residue mutations on PAR4-stimulated G protein signalling

β-arrestin-mediated desensitization of GPCR G protein signalling is a well-established phenomeon^38–41^. Within the PAR family of receptors, it has been demonstrated that PAR2 phosphorylation is a requirement for β-arrestin recruitment, ultimately leading to desensitization of the receptor ^42,43^. In keeping with the well-established role of β-arrestins in GPCR desensitization, we previously demonstrated that of β-arrestin-1/-2 CRISPR/Cas9 knockout HEK-293 cells had both increased and prolonged calcium signalling following PAR2 stimulation ^21^. Previously, it was reported that serine/threonine CT mutations in PAR1 increased G protein signalling, while analogous mutations to PAR4 had no impact on signalling^44^. To determine contribution of CT phosphorylation patterns on β-arrestin-mediated PAR4 receptor desensitization, we recorded calcium signalling in cells expressing two phosphorylation-site mutant receptors. PAR4^T363A/S366A/S369A^-YFP-mediated calcium signalling (EC_50_ = 14.0 ± 4.1 µM, Max. = 23.7 ± 3.3% A23187) was not significantly different than that elicited by the wild-type PAR4-YFP receptor (EC_50_ = 21.1 ± 3.5 µM, Max. = 29.6 ± 1.0% A23187) in response to AYPGKF-NH_2_ stimulation (Fig. 6A). Similarly, the EC_50_ of thrombin-stimulated calcium signalling from PAR4^T363A/S366A/S369A^-YFP (EC_50_ = 0.8 ± 0.2 units/mL) was comparable to PAR4-YFP (EC_50_ = 0.5 ± 0.2 units/mL). When Gα_13_ signalling was assessed we observed a significant decrease in the net BRET recorded at the highest concentrations in response to both AYPGKF-NH_2_ [EC_50_ = 14.6 ± 4.7 µM, Max. = -0.11 ± 0.01 net BRET (p < 0.0001), net BRET at 300 µM = -0.12 ± 0.01 (p < 0.05), Fig. 6B] or thrombin [EC_50_ = 2.7 ± 0.9 units/mL, Max. = -0.11 ± 0.01 net BRET, net BRET at 300 µM = -0.09 ± 0.01 (p < 0.05), Fig. 6H]. Unlike Gα_q/11_ and Gα_13_ activation, we observed no significant differences in Gα_oB_ activation with either agonist tested in PAR4^T363A/S366A/S369A^-YFP signalling compared to wild-type [AYPGKF-NH_2_ -EC_50_ = 66.0 ± 35.8 µM, Max. = -0.15 ± 0.03 net BRET, net BRET at 300 µM = -0.13 ± 0.01, Fig. 6C; Thrombin - EC_50_ = 3.7 ± 2.9 units/mL, Max. = -0.13 ± 0.04 net BRET, net BRET at 300 µM = -0.09 ± 0.01, Fig. 6I].

**Fig. 6.**
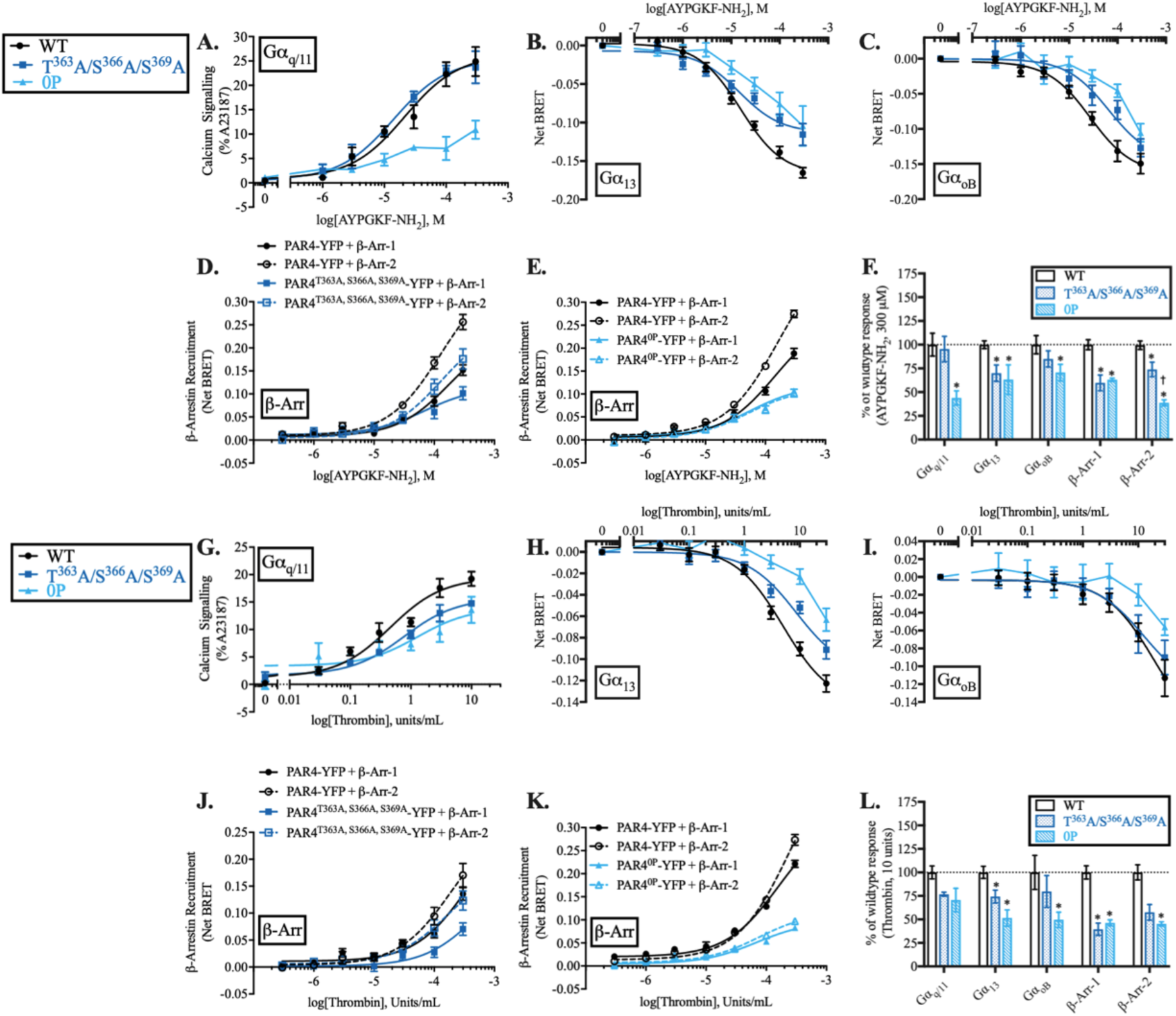
CT phosphorylation barcode and phosphorylatable residue mutations decrease β-arrestin-1/-2 recruitment to both AYPGKF-NH_2_- and thrombin-stimulated PAR4 and phospho-null PAR4 CT mutation impacts appropriate G protein activation. *G protein activation (A-C, H-J).* Calcium signalling from wild-type PAR4-YFP, PAR4^T363A/S366A/S369A^-YFP, and phospho-site null PAR4-YFP (PAR4^0P^-YFP) receptors in response to AYPGKF-NH_2_ (A) or thrombin (G) stimulation. Nonlinear regression curve fits are shown (mean ± S.E.) for three to four independent experiments (*n = 3-4*). G protein activation (TRUPATH) was recorded in cells expressing wild-type or mutant PAR4-YFP receptors in response to either AYPGKF-NH_2_ [HEK-293 cells; Gα_13_ (B), Gα_oB_ (C)] or thrombin [PAR1-KO-HEK-293 cells; Gα_13_ (H), Gα_oB_ (I)]. Nonlinear regression curve fits are shown (mean ± S.E.). Technical replicates were collected in triplicate for each experimental replicate (*n = 3-4*). β*-arrestin recruitment (D-E, J-K).* Recruitment of β-arrestins to wild-type or mutant PAR4-YFP receptors following stimulation with AYPGKF- NH_2_ [β-Arr-1 (D), β-Arr-2 (E)] or thrombin [β-Arr-1 (J), β-Arr-2 (K)]. Nonlinear regression curve fits are shown (mean ± S.E.) with triplicate data points for each concentration and receptor collected within each experiment. (*n = 3-4)*. *Summary data normalized to wild-type receptor (F, L)*. For ease of comparison, signalling of mutant PAR4 receptors was normalized to signalling recorded from wild-type PAR4 receptor in response to AYPGKF-NH_2_ (F) or thrombin (L) at the highest concentrations tested (300 µM AYPGKF-NH_2_, 10 units/mL thrombin) for each of the G protein activation and β-arrestin recruitment assays (A-E, G K). Significance was calculated on raw (non-normalized) values for each assay by T-test and is indicated (*p < 0.05 compared to wild-type PAR4-YFP receptor).

When calcium signalling was recorded in the phosphorylation-site-null PAR4^0P^-YFP receptor, peptide-stimulated calcium signalling was significantly decreased [EC_50_ = 20.9 ± 15.3 µM, Max. = 10.3 ± 1.4% A23187, Signal at 300 µM = 10.9 ± 1.9 (p < 0.05), Fig. 6A] compared to wild-type receptor at 300 µM (EC_50_ = 21.4 ± 7.7 µM, Max. = 26.3 ± 2.1% A23187, Fig. 6A). Interestingly, thrombin-stimulated signalling was unaffected (EC_50_ = 1.2 ± 1.1 units/mL, Max. = 13.8 ± 2.6% A23187, Signal at 10 units/mL = 13.6 ± 2.4% A23187, Fig. 6G) compared to wild-type receptor (EC_50_ = 0.5 ± 0.1 units/mL, Max. = 19.4 ± 2.6%; Fig. 6G). Therefore, unlike PAR2 which had significantly increased and prolonged calcium signalling in the absence of β-arrestins, removal of PAR4-phosphorylation sites does not significantly enhance calcium signalling, indeed it impairs Gα_q/11_ coupled calcium signaling in response to a peptide agonist.

We were surprised to observe that, in addition to the decreased signalling observed with AYPGKF-NH_2_-stimulated Gα_q/11_ signalling, both Gα_13_ and Gα_oB_ signalling was also decreased in response to either AYPGKF-NH_2_ [Gα_13_ – EC_50_ = *n.d.*, Max. = *n.d.*, net BRET at 300 µM = -0.10 ± 0.03 (p < 0.05), Fig. 6B; Gα_oB_ – EC_50_ = *n.d.*, Max. = *n.d.*, net BRET at 300 µM = -0.11 ± 0.01 (p < 0.05), Fig. 6C] or thrombin [Gα_13_ – EC_50_ = *n.d.*, Max. = *n.d.*, net BRET at 10 units/mL = - 0.06 ± 0.01 (p < 0.05), Fig. 6H; Gα_oB_ – EC_50_ = *n.d.*, Max. = *n.d.*, net BRET at 10 units/mL= -0.06 ± 0.01 (p < 0.05), Fig. 6I] stimulation of PAR4^0P^-YFP compared to the wild-type receptor.

### Role for phosphorylation barcode motif and CT phosphorylation sites in differential recruitment of β-arrestins to PAR4 dependent on either AYPGKF-NH_2_ or thrombin-revealed tethered ligand

A role for phosphorylation-dependent GPCR-mediated recruitment of β-arrestins is well established^45,46^. The importance of phosphorylation codes/motifs in β-arrestin binding and signalling has also been indicated for many GPCRs including the β2-adrenergic receptor, rhodopsin receptor and vasopressin receptors ^47–50^. A so-called “complete” phosphorylation barcode motif, involves three phosphorylatable residues interspaced by two residues [PxxxPxxP, wherein “P” denotes a phosphorylatable serine or threonine residue, and “x” denotes any other residue], which have been shown to be a key component of receptor CT/arrestin interaction ^51^. Additionally, partial barcode motifs have been identified wherein a phosphorylatable residue spot within the motif is occupied by another residue, frequently an acidic residue ^51^. Phosphorylation motifs in the PAR4 C-terminal tail were identified using the PhosCoFinder tool ^51^ which identified one complete phosphorylation motif -Thr^363^/Ser^366^/Ser^369^. Mutation of this motif to alanine (PAR4^T363A, S366A, S369A^-YFP) significantly reduced β-arrestin recruitment to PAR4 in response to AYPGKF-NH_2_ stimulation of the receptor [β-Arr-1 Max. = 0.10 ± 0.01 net BRET (p < 0.05), β-Arr-2 Max. = 0.18 ± 0.02 net BRET (p < 0.05), Fig. 6D) compared to wild-type receptor (β-Arr-1 Max. = 0.15 ± 0.01 net BRET, β-Arr-2 Max. = 0.26 ± 0.02 net BRET, Fig. 6D).

Interestingly, in response to thrombin stimulation only β-arrestin-1 recruitment was significantly decreased [Max. = 0.07 ± 0.01 net BRET (p < 0.05), Fig. 6J] compared to wild-type receptor (Max. = 0.14 ± 0.01 net BRET, Fig. 6J); while β-arrestin-2 recruitment [Max. = 0.12 ± 0.02 net BRET (p = 0.11), Fig. 6J] was not statistically different than thrombin-stimulated recruitment to wild-type receptor (Max. = 0.17 ± 0.02 net BRET, Fig. 6J). Our data therefore implicate the CT barcode motif Thr^363^/Ser^366^/Ser^369^ in peptide-stimulated recruitment of β-arrestin-1 and -2 to PAR4 and thrombin-stimulated β-arrestin-1 recruitment.

Given that we do not observe a complete loss of β-arrestin-1/-2 recruitment with the complete barcode motif mutation (Thr^363^Ala/Ser^366^Ala/Ser^369^Ala), we generated a mutant PAR4 receptor wherein all of the CT serine/threonine residues are mutated to alanine (PAR4^0P^-YFP) to determine if there was any further loss of recruitment. Recruitment of β-arrestin-1 to PAR4^0P^-YFP was significantly reduced compared to wild-type receptor, however, was not significantly more detrimental than the barcode motif mutation alone [AYPGKF-NH_2_ -β-Arr-1 Max. = 0.11 ± 0.00 net BRET (p < 0.0001 compared to wild-type; p = 0.70 compared to PAR4^T363A, S366A, S369A^-YFP), Fig. 6E; thrombin - β-Arr-1 Max. = 0.08 ± 0.01 net BRET, (p < 0.0001 compared to wild-type; p = 0.35 compared to PAR4^T363A, S366A, S369A^-YFP), Fig. 6K] (Fig. 6F & 6L). Additionally, β-arrestin-2 recruitment in response to thrombin stimulation of PAR4 was significantly decreased compared to wild-type receptor, however, not more so than the barcode mutation alone [β-Arr-2 Max. = 0.10 ± 0.00 (p < 0.0001 compared to wild-type; p = 0.23 compared to PAR4^T363A/S366A/S369A^-YFP), Fig. 6K & 6L]. Interestingly, we observed a further reduction of β-arrestin-2 recruitment to PAR4 in response to peptide stimulation in PAR4^0P^-YFP compared to barcode motif mutation alone [β-Arr-2 Max. = 0.10 ± 0.01 (p < 0.0001 compared to wild-type; p < 0.05 compared to PAR4^T363A/S366A/S369A^-YFP, Fig. 6E & 6F). These data may therefore highlight a role for agonist-dependent subtype selectivity, as was observed with the mutation of the complete phosphorylation barcode and reveal residues involved in arrestin-subtype selectivity in the CT of PAR4. These data, however, should be carefully followed up with further structural and functional studies to identify specific residues that enable this effect.

### Alphafold 3 predicted interactions between PAR4 and signaling effectors

Finally we modeled PAR4 interactions with β-arrestin-1/-2, Gα_q_, Gα_11_, Gα_12_, Gα_13_, Gα_i1_, and Gα_o_ using AlphaFold 3 (Fig. 7). The models revealed multiple contacts between β-arrestins and the phosphorylated C-tail of PAR4. Additionally, β-arrestins formed contacts with PAR4 intracellular loops and transmembrane domains, suggesting the capture of a core conformation. While the models showed interaction between PAR4 H8 residue (Ser^8.47^) and β-arrestin-1, no H8 interactions were observed with β-arrestin-2. The models also predicted several interactions between PAR4 H8 and CT residues with Gα_q_. However, for Gα_11_, Gα_i1_, Gα_o_, and Gα_12/13_, the predicted interactions were predominantly confined to the intracellular loops and transmembrane domain 6. The H8 residues identified through mutagenesis as critical for PAR4 interaction with signaling effectors may therefore mediate intermediate steps in these interactions that are not fully captured by our modeling approach. Further functional or structural studies will be needed to validate these predicted interactions.

**Figure 7.**
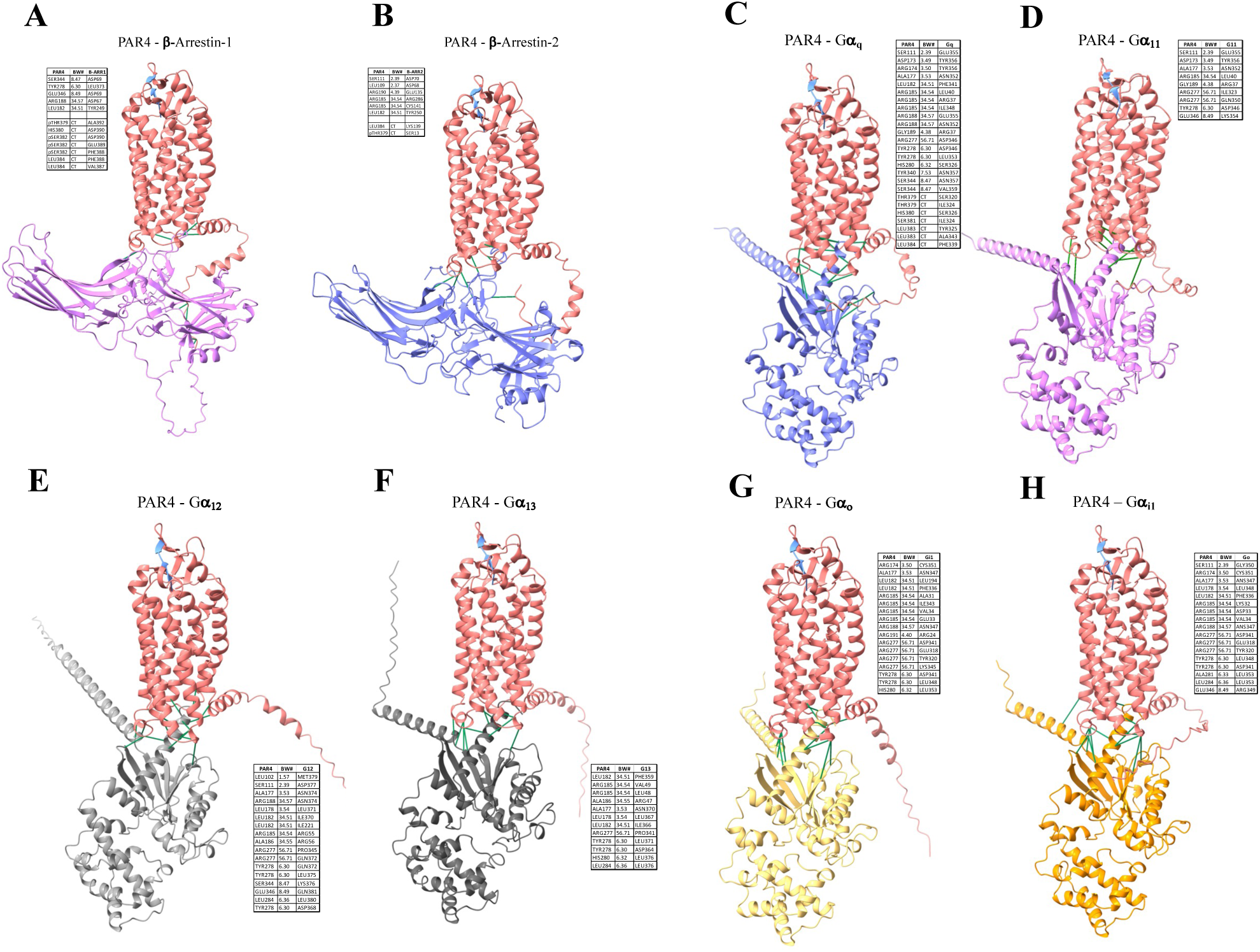
Alphafold3 prediction of PAR4 interaction with β-arrestins and G proteins. Tethered-ligand exposed PAR4 interactions with β-arrestin-1/-2, Gα_q_, Gα_11_, Gα_12_, Gα_13_, Gα_o_ and Gα_i1_ were modeled using alphafold3. β-arrestin interactions were modeled with phosphorylated C-tail serine and threonine residues while G proteins were modeled without any post-translational modifications. Contacts within 3.5Å were identified and contacts are depicted as green pseudobonds and listed in the adjacent tables. In all cases the exposed tethered-ligand, docked in the orthosteric ligand binding pocket is depicted in blue. Ballesteros-Weinstein numbering (BW#) is from GPCRdb.

## Discussion

Here we examined residues found within the H8 and CT as determinants of PAR4 coupling to several major signalling effectors including Gα_q/11_, Gα_12/13_, Gα_i/o_, and β-arrestin. We find that residues in the PAR4 H8 (Lys^350^, Leu^355^, Phe^356^, Gln^357^) are necessary for appropriate G protein activation and signalling, as well as β-arrestin-1/-2 recruitment. Additionally, we find that a TM7/H8 interaction (Tyr^340^/Phe^347^) is important for receptor signalling activity. Further, we demonstrate that a phosphorylation barcode (Thr^363^/Ser^366^/Ser^369^) is crucial for β-arrestin-1/-2 recruitment to PAR4. Interestingly, we also observed key differences in the relative importance of these sites, dependent on whether PAR4 was activated by the proteolytically-revealed tethered ligand or through agonist peptide stimulation. Thus, this study highlights the importance of H8 and CT residues in PAR4 signalling as well as underscores the subtle differences in effector coupling triggered by different modes of receptor activation.

### H8 R^352^AGLFQRS^359^ Motif

The H8 and C-terminal tail of GPCRs are key sites of recruitment, interaction, and activation of intracellular effectors, such as G proteins and β-arrestins. Structures of Gα_s_-, Gα_i_-, Gα_o_-, and Gα_q_-bound GPCRs show that residues within the H8 of adenosine 2A (A_2A_R), β_2_AR, µ-opioid (MOR), adenosine A1 (A_1_R), serotonin 1B (5HT_1B_R), rhodopsin, and 5-HT_2A_ receptors make contacts with the α and β (β1 or β2) subunits of their respective G proteins ^52,53^. Conversely, studies of dopamine D1 and D2 receptor H8 chimeras have revealed that the H8 is not involved in dopamine receptor G protein coupling but is important for β-arrestin-mediated receptor desensitization of these receptors ^54^. Previously, we demonstrated that deletion of an eight amino acid sequence in the H8 of PAR4 (R^352^AGLFQRS^359^) abrogates Gα_q/11_-mediated calcium signalling and significantly decreased β-arrestin-1/-2 recruitment ^30^. In the current study we observed that mutation of this entire sequence to alanine (PAR4^R352-S359A^-YFP) mimicked the abrogation of calcium signalling and reduced β-arrestin recruitment previously reported with the deletion mutant.

Residues within the PAR1 H8 (S^375^SECQRYVYSILCC^388^) were also shown to be necessary for Gα_q_ signalling. Specifically, mutation of hydrophilic residues (Gln^379^Ala/Arg^380^Ala) significantly reduced Gα_q_-PLC-β-dependent inositol phosphate production by approximately 25% in response to both thrombin and peptide activation of PAR1^55^. When this mutation was combined with intracellular loop 1 mutant, Lys^135^Ala, a 40-50% reduction was observed in Gα_q_-PLC-β activity, highlighting the importance of an ICL1/H8 activation motif in PAR1-mediated Gα_q_ activation^55^. Mutational studies of the rhodopsin receptor revealed a different mechanism wherein Arg^314^/Asn^315^ mutation, located at analogous sites to PAR1 Gln^379^/Arg^380^ (8.51/8.52), had no impact on Gα_t_ activation; however, mutation of hydrophobic residues (Phe^313^, Met^8.54^) significantly reduced Gα_t_ activation ^26,56^. Interestingly, mutation of the analogous PAR1 residues (Cys^378^, Val^382^) did not impact Gα_q_-coupling suggesting a role for H8 in G protein subtype selectivity^55^. Similar to findings with rhodopsin, mutation of hydrophobic residues (Leu^404^, Phe^408^, and Phe^412^) in the H8 of cannabinoid 1 receptor (CB_1_) decreased G protein activation, while mutation of basic residue had no effect^57^. Here we identified that H8 residues Leu^355^-Gln^357^regulated PAR4 G protein activation. Interestingly, we observed that mutation of these residues impacts on both peptide- and enzyme-activated PAR4 mediated Gα_q/11_-coupled calcium signalling. Therefore PAR4, like PAR1, requires a H8 glutamine for efficient Gα_q/11_ activation. Like in CB_1_, mutation of leucine may remove a necessary hydrophobic residue for G protein binding and activation^57^.

Following a similar strategy, we investigated β-arrestin recruitment to the segmental mutations of the R^352^AGLFQRS^359^ motif. As with calcium signalling, β-arrestin-1/-2 recruitment was significantly reduced with PAR4 Leu^355^-Gln^357^Ala mutation in response to both thrombin and peptide stimulation. Both peptide- and thrombin-stimulated β-arrestin-1 recruitment to PAR4 was decreased with Arg^358^-Ser^359^Ala mutation however, only peptide-stimulated β-arrestin-2 recruitment was diminished in this mutant. Further, we observed a reduction in thrombin-stimulated β-arrestin-2 recruitment to PAR4 with Arg^352^-Gly^354^Ala mutation. Interestingly, Arg^352^-Gly^354^Ala mutation enhanced both β-arrestin-1 and -2 recruitment following peptide stimulation. Thus, the overall decrease observed with Leu^355^-Gln^357^Ala mutation may represent a global phenotype since this mutation also decreased PAR4 calcium signalling, but there may be a role for other residues within the R^352^AGLFQRS^359^ motif in arrestin-subtype selectivity and agonist-dependent differences in modulation of β-arrestin-1/-2 recruitment.

Residues within the H8 has been implicated in regulating β-arrestin recruitment to other GPCRs. For example H8 domain swap experiments in the dopamine D1 (D1R) and D2 (D2R) receptors revealed a role for H8 in β-arrestin-1/-2-mediated desensitization in D1R, but not D2R^54^. The H8 of the rhodopsin receptor has also been implicated in mediating phosphate sensing through changes in conformational dynamics during the “pre-binding” state preceding CT phosphorylated residues interaction with the polar core of visual arrestin^58^. Therefore, our findings here should be followed up with studies evaluating whether mutations perturb interactions of the PAR4 H8 with β-arrestins directly or whether mutations in H8 impact conformational dynamics of H8 resulting in the diminished β-arrestin recruitment observed.

The H8 of many GPCRs are known to interact with other regions of the GPCR or the plasma membrane to stabilize receptor conformations which are crucial to maintain both inactive and active conformations as well as mediate effector interaction^36,55,58–60^. In our current investigation of the H8 R^352^AGLFQRS^359^ sequence we have not extensively evaluated the potential loss of intramolecular contacts with other regions of the receptor and this remains an area for future investigation.

### H8 Lys^350^ and CT Lys^367^

The PAR4 CT contains two lysine residues – Lys^350^ in the H8 and Lys^367^ distal to H8. Mutation of Lys^350^ to alanine significantly decreased PAR4-mediated Gα_q/11_, Gα_13_, and Gα_oB_ activation and β-arrestin-1/-2 recruitment following peptide stimulation and Gα_q/11_ and Gα_oB_ with thrombin activation of PAR4. Interestingly, Lys^367^Ala mutation also significantly reduced peptide-stimulated calcium signalling from PAR4, however, not as significantly as H8 Lys^350^Ala mutation and with no detrimental impact on peptide- or thrombin-stimulated β-arrestin recruitment. While the mechanism underlying the contribution of Lys^350^ to Gα_q/11_, Gα13, and Gα_oB_ binding were not investigated in this study, there is a clear and supported role for charged H8 lysine residue in direct G protein interactions demonstrated in other class A GPCRs. Mutational studies with the muscarinic 3 receptor (M3R) revealed lysine within the N-terminal domain of H8 made contacts with the α4/β6 loop of Gα_q/11_^35^. Further, studies with µ-opioid receptor (µOR) implicate H8 lysine as having decreased solvent accessibility upon G protein binding following activation^33^. Similar to our observations with PAR4 Lys^350^Ala mutation (8.53), Lys^320^Gln (8.52) mutation in MCH_1_R decreased MCH-stimulated, Gα_q_-mediated calcium mobilization ^24^. It is important to note that H8 lysine mutations in several GPCRs, including the melanin-concentrating hormone (MCH) receptor 1 (MCH_1_R) and bradykinin B_2_ receptor (B_2_R), resulted in reduced plasma membrane expression^24,25^. To ensure that reduction in G protein activation was not due to poor membrane localization of lysine mutant PAR4 receptor, we examined and observed by confocal microscopy that PAR4 Lys^350^Ala, Lys^367^Ala, and Lys^350^Ala/Lys^367^Ala double mutant expressed appropriately on the cell membrane.

As previously stated, mutation of Lys^350^ decreased both peptide- and thrombin-stimulated β-arrestin-1/-2 recruitment, while Lys^367^Ala had no effect on agonist-stimulated recruitment. Additionally, combined mutation was no more detrimental than Lys^350^Ala mutation alone. Previous studies of the thyrotropin-releasing hormone (TRH) receptor (TRH_1_R) and B_2_R revealed that H8 lysine residues were important in mediating GRK interaction and CT phosphorylation of the receptors. Mutation of TRH_1_R H8 Lys^326^ was found to significantly reduce CT phosphorylation and receptor internalization, which could be partially overcome with overexpression of G protein receptor kinase 2 (GRK2)^61^. Similarly, H8 lysine mutation to proline in B_2_R (Lys^315/8.53^; analogous to Lys^350/8.53^ residue in PAR4) decreased agonist-stimulated CT phosphorylation and agonist-dependent internalization, which could be recovered in part by overexpression of GRK2 or GRK3^25^. Thus, alterations in GRK-mediated receptor phosphorylation may be responsible for the reductions observed with Lys^350^ mutation and should be investigated in future studies.

### DPxxYx_6_F motif

Many Class A GPCRs possess a canonical NPxxYX_5,6_F motif/domain, spanning TM7 and H8, including Asn^7.49^, Pro^7.50^, Tyr^7.53^, and H8 Phe^8.50^. This motif is involved in stabilizing the inactive state of Class A GPCRs. In crystal structures of inactive state GPCRs, the side chain of Tyr^7.53^ points towards helices I, II, or VIII, whereas, structures of the active state have Tyr^7.53^ changing rotamer conformation to face the interior of the transmembrane bundle, pointing towards helices VI and III^62^. Additionally, an inactive-state stabilizing water molecule network involving the TM7 NPxxY, TM3 E/DRY, and TM6 WXPF/Y motifs may be partially constituted by Tyr^7.53 37^. Previous studies of rhodopsin and the β_2_-adrenergic receptor have demonstrated that mutations in this motif alter ligand affinity, receptor plasma membrane expression, G protein coupling, and interactions with small G proteins, such as RhoA, dependent on the receptor studied ^26–29^. In our mutational study, we observed a significant loss of Gα_q/11_, Gα_13_, and Gα_oB_ signalling with mutation of Tyr^340^-Ala and of Gα_q/11_ and Gα_oB_ with Phe347-Ala mutations, irrespective of the stimulating PAR4-activating agonist. In all cases there was no impact on appropriate cell membrane localization (Supplementary Fig. 1) of the receptor.

Functional studies have highlighted that the absence of side-chain interaction between Tyr^7.53^ and Phe^8.50^ increases the population of active rhodopsin (Meta II state) with no concomitant increase in transducin (Gα_t_) activation^26,37^. Further, these studies show that hydrophobic side chain interaction between Tyr^7.53^ and Phe^8.50^ constitutes a constraint that becomes necessarily perturbed during receptor interaction to allow for conformational rearrangement of H8 and G protein binding^26,37^. Mutation of rhodopsin H8 Phe^313^ (8.50) to alanine caused a corresponding loss of Gα_t_ activation with rhodopsin^26^. Further, perturbation of this interaction, as well an interaction between TM3 3.46 and TM6 6.37 residues, enables Tyr^7.53^ to make a new contact with TM6 6.37 residue and form an interaction with the α5 helix of the G protein, which has been demonstrated in several Class A GPCRs (rhodopsin, M2R, µOR, β_2_AR, and A_2A_R)^63^. Mutation of Tyr^7.53^ in the V2 vasopressin receptor (V2R) also significantly reduces the receptor-mediated activation of both Gα_s_ and Gα_q_ ^63^. While it is not possible in this study to resolve if a change in the active-state population of PAR4 or a perturbation in the conformational rearrangement of TM7/H8 interaction is responsible for the loss of signalling observed, it is clear that these residues, especially Tyr^340^, have a role in Gα_q/11_, Gα_13_, and Gα_oB_ signalling downstream of PAR4 activation.

In addition to substantial reductions in G protein activation, we observed significant decreases in both thrombin- and peptide-stimulated recruitment of β-arrestin-1/-2 to PAR4 DPxxYx_6_F mutants, Tyr^340^Ala and Phe^347^Ala. The NPxxYx_5,6_F motif has been shown for several Class A GPCRs to be involved in interactions with β-arrestins. Mutations of this motif are thought to perturb GRK interactions with the GPCR and thus decrease GRK-mediated phosphorylation^33–35^. In studies of the β_2_-adrenergic receptor (β_2_AR), mutation of TM7 Tyr^326^ (7.53) decreased GRK-mediated phosphorylation and receptor sequestration, which was reversible through overexpression of GRKs 2-6 ^27,64,65^. Similar to our observation in the PAR4 Tyr^340^Ala and Phe^347^Ala mutations, β-arrestin recruitment is significantly reduced in both β_2_AR and α_1B_AR receptors with TM7 Tyr^7.53^ mutations^66^. Whether changes in GRK-mediated receptor phosphorylation of PAR4 accounts for the reduction in β-arrestin recruitment observed with Tyr^340^Ala and Phe^347^Ala mutations should be the focus of further study.

### Phosphorylation-site mutations

A role for phosphorylation-dependent GPCR-mediated recruitment of β-arrestins is well established^45,46^. The importance of a phosphorylation barcode motif in β-arrestin binding and signalling has also been demonstrated for many GPCRs including the β2-adrenergic receptor, rhodopsin receptor, chemokine receptors 4 & 7, vasopressin, and angiotensin II receptor type 1 receptors^47–50^. It has been reported that 52.3% of Rhodopsin family GPCRs possess either a full or partial phosphorylation barcode^51^. These GPCRs can be further divided into two classes based on their β-arrestin-1 and -2 binding characteristics. Class A GPCRs bind β-arrestin-2 more tightly than β-arrestin-1 and are typified by receptors such as α1- and β2-adrenergic, μ-opioid, and dopamine D_1_A receptors. Class B GPCRs bind β-arrestins-1/-2 equally well (e.g. angiotensin II type 1A receptor, neurotensin receptor 1, vasopressin-2 receptor, thyrotropin-releasing hormone receptor, and substance P receptor)^67^. All class A β-arrestin-binding GPCRs have, at most, one complete phosphorylation barcode (PxxPxxP, as previously defined)^51^. Interestingly, our data consistently show more robust recruitment of β-arrestin-2 versus β-arrestin-1 and the PAR4 CT possesses only one such complete phosphorylation barcode. Thus, PAR4 may be considered a class A β-arrestin-binding GPCR. Further, when the complete phosphorylation barcode is mutated (Thr^363^Ala/Ser^366^Ala/Ser^369^Ala), we observe significantly reduced β-arresin-1/-2 recruitment in response to both peptide and thrombin activation of the receptor compared to wild-type receptor.

While some phosphorylation barcodes may contribute to β-arrestin recruitment to activated GPCRs, other phosphorylated residues may play a role in β-arrestin subtype-specificity, the recruitment of other downstream signalling partner (such as SNX27 for β_2_AR) and mediate of affinity of β-arrestin binding to GPCRs^47,48,51,68^. We therefore evaluated β-arrestin recruitment to a CT phospho-null PAR4 receptor. Interestingly, only peptide-stimulated β-arrestin-2 recruitment was further reduced compared to Thr^363^Ala/Ser^366^Ala/Ser^369^Ala mutation alone. Thus, these data support a predominant role for the complete phosphorylation barcode in β-arrestin recruitment to activated PAR4.

When we evaluate the impact of reduced β-arrestin-1/-2 recruitment to PAR4 on G protein signalling, we observed no significant increase or prolongation in agonist-stimulated calcium signalling compared to wild-type receptor. This is in stark contrast to what we previously observed with PAR2-mediated signalling wherein, maximal calcium signalling increases and demonstrates prolonged kinetics compared to conditions when β-arrestin recruitment is impaired^21^. These data are in agreement with previously published mutational studies of serine/threonine residues in the PAR4 CT which had no effect on increasing PAR4 G protein signalling^44^. This suggests that while β-arrestin recruitment is reduced in the PAR4 phosphorylation barcode motif mutant, the level of recruitment may be sufficient to desensitize the calcium signalling pathway. Alternatively, there may be other mechanisms enabling PAR4 desensitization such as ubiquitination and internalization as observed with PAR1 and PAR2^69^. Intriguingly, we observed decreased activation of G protein signalling in the phosphorylation null PAR4 mutant (but not phosphorylation barcode mutated PAR4) which suggest that one or more of these CT residues may serve a functional role in G protein activation, the mechanisms of which are not readily apparent but may involve direct interactions and warrant future investigation.

## Conclusion

Overall, our findings unveil crucial molecular mechanisms governing PAR4-stimulated G protein and β-arrestin-1/-2 signalling and regulation. Further, these data highlight important divergences in the relative importance of H8 and CT residues on PAR4 signalling, dependent on peptide- or enzyme revealed tethered-ligand activation of the receptor. Differential activation of PAR4 by different enzymes may similarly exhibit differential regulation of effector interactions with PAR4. Alphafold 3 modelling further predicted additional interactions that may govern stable interactions between PAR4 and signaling effectors. Ultimately, understanding the molecular mechanisms governing effector interactions, signalling, and signal regulation at PAR4 provides novel insights into how this receptor functions and may aid in the development of novel therapeutics targeting PAR4.

## Materials and Methods

### Chemicals and Other Reagents

Thrombin from human plasma (catalogue no. 605195) was purchased from Millipore-Sigma (St. Louis, MI). AYPGKF-NH_2_ (> 95% purity by HPLC/MS) was purchased from Genscript (Piscataway, NJ). Agonists were prepared in 4-(2-hydroxyethyl)-1- piperazineethanesulfonic acid (HEPES, 25 mM). All other chemicals or reagents were purchased from Millipore-Sigma, Thermo Fisher Scientific, or BioShop Canada, Inc. (Burlington, Ontario, Canada), unless otherwise stated.

### Molecular Cloning and Constructs

The plasmid encoding the human PAR4 receptor with an in frame enhanced yellow fluorescent protein fusion tag (PAR4-YFP) has been previously described ^30^. eYFP fused PAR4 has been utilized in several previous studies where we have demonstrated that PAR4-YFP couples to the known PAR4 signalling pathways appropriately ^13,17,17,30^. QuikChange XL Multi Site-Directed Mutagenesis kit (Agilent Technologies, Mississauga, ON, Canada) was used to generate all H8 and the C-terminal tail PAR4 mutants described in this study. Additionally, wild-type and mutant PAR4 constructs were generated with C-terminal HA tags (2x HA-Stop) for use in TRUPATH signalling assays. All constructs were verified by Sanger sequencing (London Regional Genomics Centre, University of Western Ontario).

### Cell Lines and Culture Conditions

All media and cell culture reagents were purchased from Thermo Fisher Scientific (Waltham, MA). Human embryonic kidney (HEK) cells (HEK-293; ATCC) and PAR1-knockout HEK-293 ^70^ cell lines were maintained in Dulbecco’s modified Eagle’s medium supplemented with 10% fetal bovine serum, 1% sodium pyruvate, and penicillin streptomycin solution (50,000 units penicillin, 50,000 μg streptomycin). Since trypsin activates PAR4, cells were routinely sub-cultured using enzyme-free isotonic phosphate-buffered saline (PBS) containing EDTA (1 mM). Cells were transfected with PAR4-YFP or mutated PAR4-YFP receptor vectors using X-tremeGENE 9 (Millipore-Sigma). Transiently transfected cells were always assayed or imaged at 48 hours post-transfection to ensure consistent levels of protein expression.

### Calcium Signalling

Agonist-stimulated calcium signalling was recorded in HEK-293 and PAR1-knockout HEK-293 cells, as previously described ^13,71,72^. Cells were detached in enzyme-free cell dissociation buffer, pelleted, and resuspended in Fluo-4 NW (no wash) calcium indicator dye (Thermo Fisher Scientific). Following a 30-minute incubation at ambient temperature, intracellular fluorescence (excitation 480 nm; emission recorded at 530 nm) was monitored before and after addition of agonists (thrombin, PAR1-knockout HEK-293; or AYPGKF-NH_2_, HEK-293) on a PTI spectrophotometer (Photon Technology International, Birmingham, NJ). Responses were normalized to the fluorescence obtained with calcium ionophore (A23187, 3 μM; Sigma-Aldrich).

### Measurement of G protein activation with TRUPATH

G protein activation in response to thrombin or AYPGKF-NH_2_-mediated activation of PAR4 was recorded with wild-type and mutant PAR4 constructs using TRUPATH biosensors (Addgene, Kit# 1000000163) ^22^. Cells were plated to a density of 2.1-2.3 x 10^6^ cells/dish in 10 cm dishes and transfected using a 1:1:1:1 DNA ratio of receptor:Gα-RLuc8:Gβ:Gγ-GFP2 (750ng/construct), using X-tremeGENE 9 (12 µl) (Millipore-Sigma) in OptiMEM (Gibco-ThermoFisher, Waltham, MA) with the appropriate Gα-RLuc8:Gβ:Gγ-GFP2 pairings reported in Olsen et. al, 2020 ^22^. 24 hours post-transfection, cells were mechanically lifted and replated into white 96-well plates (Greiner Bio-One, Monroe, NC; Corning; Oneonta, NY). At 48 hours, cell media was removed and replaced with 60 μL of assay buffer (1x HBSS + 20 mM HEPES, pH 7.4), followed by a 10 μL addition of freshly prepared 50 μM coelenterazine 400a (Nanolight Technologies, Pinetop, AZ). After a five-minute equilibration period, cells were treated with 30 μL of drug for an additional 5 minutes. Plates were then read in an LB940 Mithras plate reader (Berthold Technologies, Oak Ridge, TN) with 395 nm (RLuc8-coelenterazine 400a) and 510 nm (GFP2) emission filters, at 1 second/well integration times. BRET2 ratios were computed as the ratio of the GFP2 emission to RLuc8 emission.

### BRET Detection of β-arrestin-1/-2 Recruitment

BRET assays were employed to detect agonist-stimulated β-arrestin-1/-2 recruitment to PAR4-YFP and mutant PAR4-YFP constructs in HEK-293 cells as described ^13,17,30^. PAR4-YFP or mutant PAR4-YFP constructs (1 µg) and Renilla luciferase-tagged β-arrestin-1 or -2 (β-arr-1 and -2-Rluc; 0.1 µg) were transiently transfected for 48 hours. Cells were plated in white 96-well culture plates (Corning; Oneonta, NY) and recruitment of β-arresitin-1/-2 to PAR4 were detected by measuring the BRET signal 20 minutes after agonist stimulation and the addition of 5 μM h-coelenterazine prior to BRET recording (NanoLight Technology, Pinetop, AZ) on a Mithras LB940 plate reader (Berthold Technologies, Bad Wildbad, Germany), as previously described ^13,30^.

### Confocal Microscopy

HEK-293 cells transiently transfected with PAR4-YFP or mutant PAR4-YFP were sub-cultured onto glass coverslips (Thermo Fisher Scientific) and analyzed by confocal microscopy to ensure appropriate cell surface expression. Cells were fixed with 4% w/v paraformaldehyde solution (methanol-free; Fisher Scientific), stained with DAPI for nuclear staining, and the subcellular receptor localization was assessed by imaging eYFP expression with an Olympus FV1000 (Centre Valley, PA).

### Alphafold3 modelling of PAR4 effector complexes

Interactions of PAR4 with signaling effectors was modelled using the Alphafold3 server (https://alphafoldserver.com). The highest ranked models were visualized in ChimeraX and contacts within 3.5Å were identified using the “alphafold contacts” command. In the case of modelling PAR4-β-arrestin-1/-2 contacts, the CT Ser and Thr residues were phosphorylated. No post-translational modifications were applied when modeling G protein interactions. The list of amino acid sequences (Table 1) and all Alphafold models and associated data are available in the supplementary information.

**Table 1.**
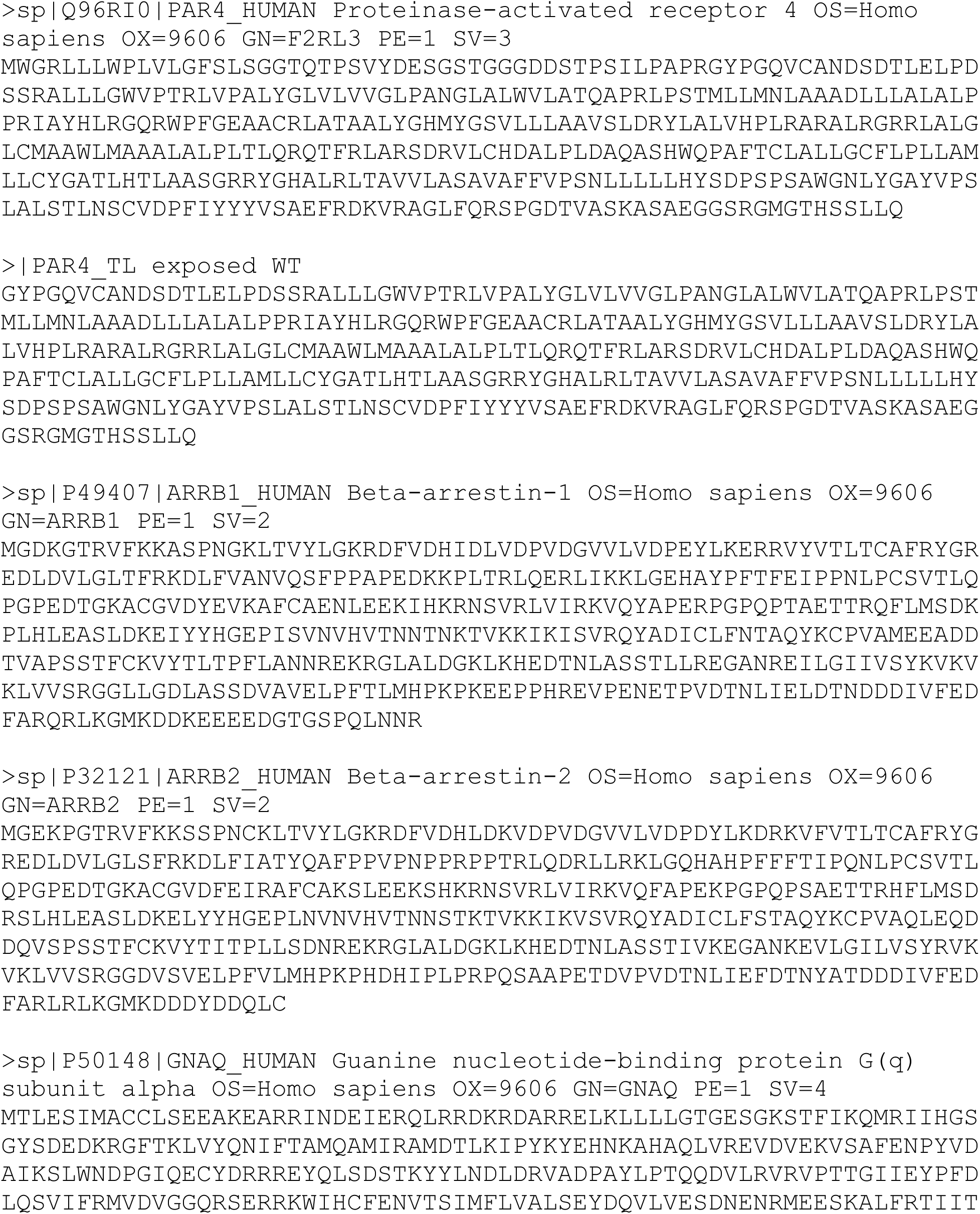
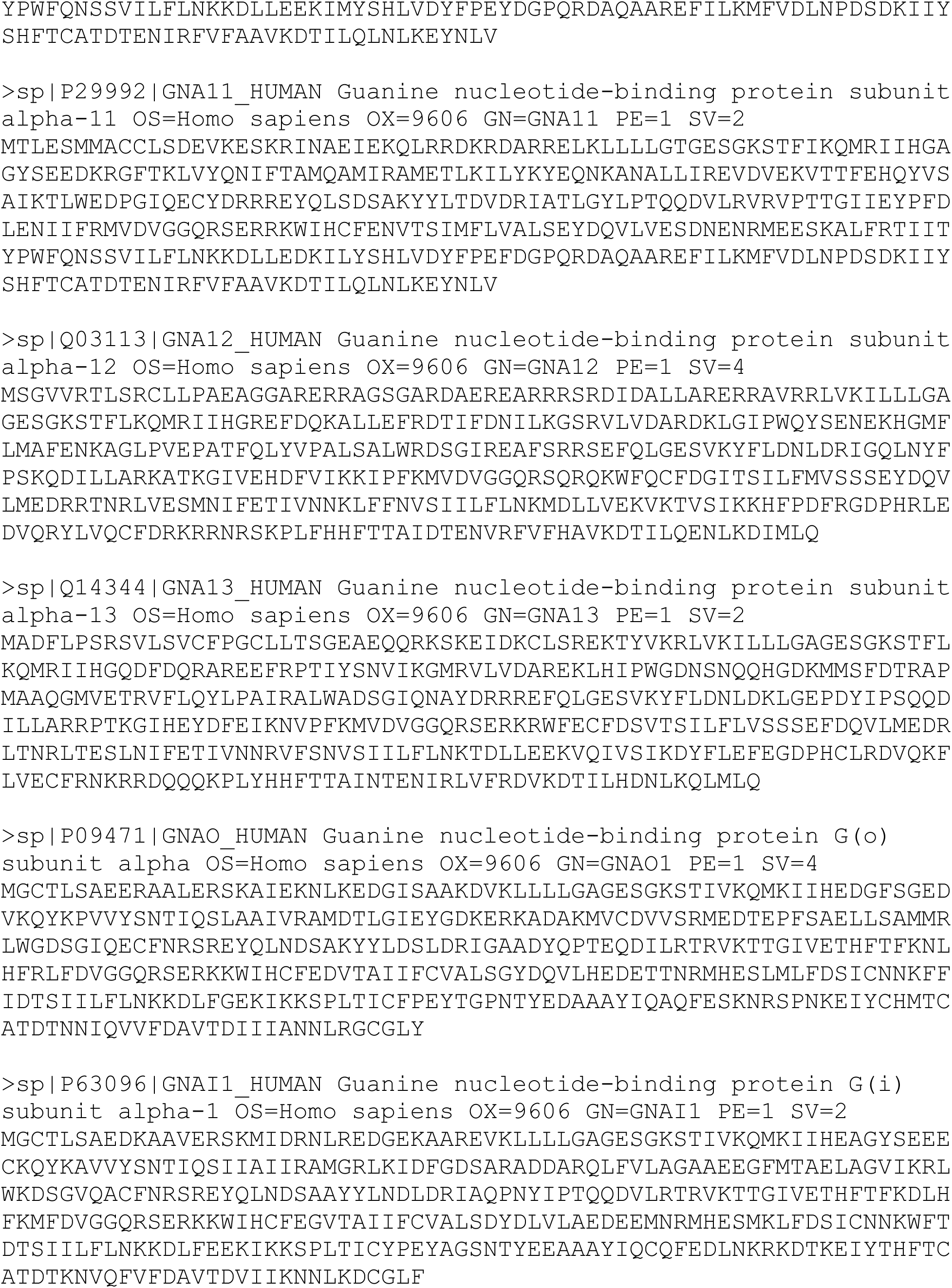
PAR4, β-arrestin and G protein sequences used for Alphafold3 prediction of protein structures.

### Statistical Analysis

Curve fitting (three-parameter, nonlinear regression) and statistical analysis was completed with Prism 7 software (GraphPad Software, La Jolla, CA). Statistical significance of EC_50_ shifts was calculated using the extra sum of squares analysis ^13,21,73^. TRUPATH BRET were conducted on all mutants and wild-type receptor simultaneously thus the PAR4-YFP curves within all Gα_13_ and Gα_oB_ graphs is repeated for reference (data and curves shown in black, circles). Net BRET, calcium signalling, and TRUPATH data obtained with AYPGKF-NH_2_ (300 µM) or thrombin (10 units/mL) were used to compare the maximal signal achieved in our study for comparison of mutant PAR4 constructs to wild-type PAR4. This has been especially important in cases where a mutant did not signal in a manner that enable robust curve-fitting for comparison. Data obtained with AYPGKF-NH_2_ (300 µM) or thrombin (10 units/mL) was normalized compared to wild-type PAR4 receptor for all mutants and is shown as a column graph within figures; however, statistical significance was calculated by T-test on the raw data values and is reported on column graphs of normalized findings for ease of the reader. Previously, we demonstrated that β-arrestin recruitment to PAR4 does not saturate upon stimulation with AYPGKF-NH_2_ and thrombin ^17,21,30^; thus, statistical significance of maximal net BRET (300 µM AYPGKF-NH_2_, 10 units/mL thrombin) was determined using t-test (*p < 0.05) and all data within the text represents the net BRET signal achieved at the highest concentration of agonists tested. Data are expressed as mean ± S.E. throughout the text, table, and figure legends.

## Acknowledgments

We thank Dr. Michel Bouvier for providing the Renilla luciferase-tagged β-arrestin-1 and -2 constructs. The TRUPATH assay kit was a kind gift from Dr. Bryan Roth.

## Data and material availability

All data supporting the findings in this study are included in the manuscript or supplementary materials. Original data or materials are available from the corresponding author upon reasonable request.

## Supplementary data

**Supplementary Figure 1.** Confocal micrographs of wild-type and mutant PAR4-YFP expression

**Supplementary Figure 2.** Thrombin stimulated PAR4 Gα_11_ TRUPATH activation

**Supplementary Figure 3.** AYPGKF-NH_2_ stimulated PAR4 Gα_11_ TRUPATH activation.

**Supplementary Table 1.** Sequences of receptor and effectors for Alphafold3 modeling.

**Supplementary data Zip folder.** PAR4 H8 and CT regulation of signaling SI AF models

**Supplementary Figure 1.**
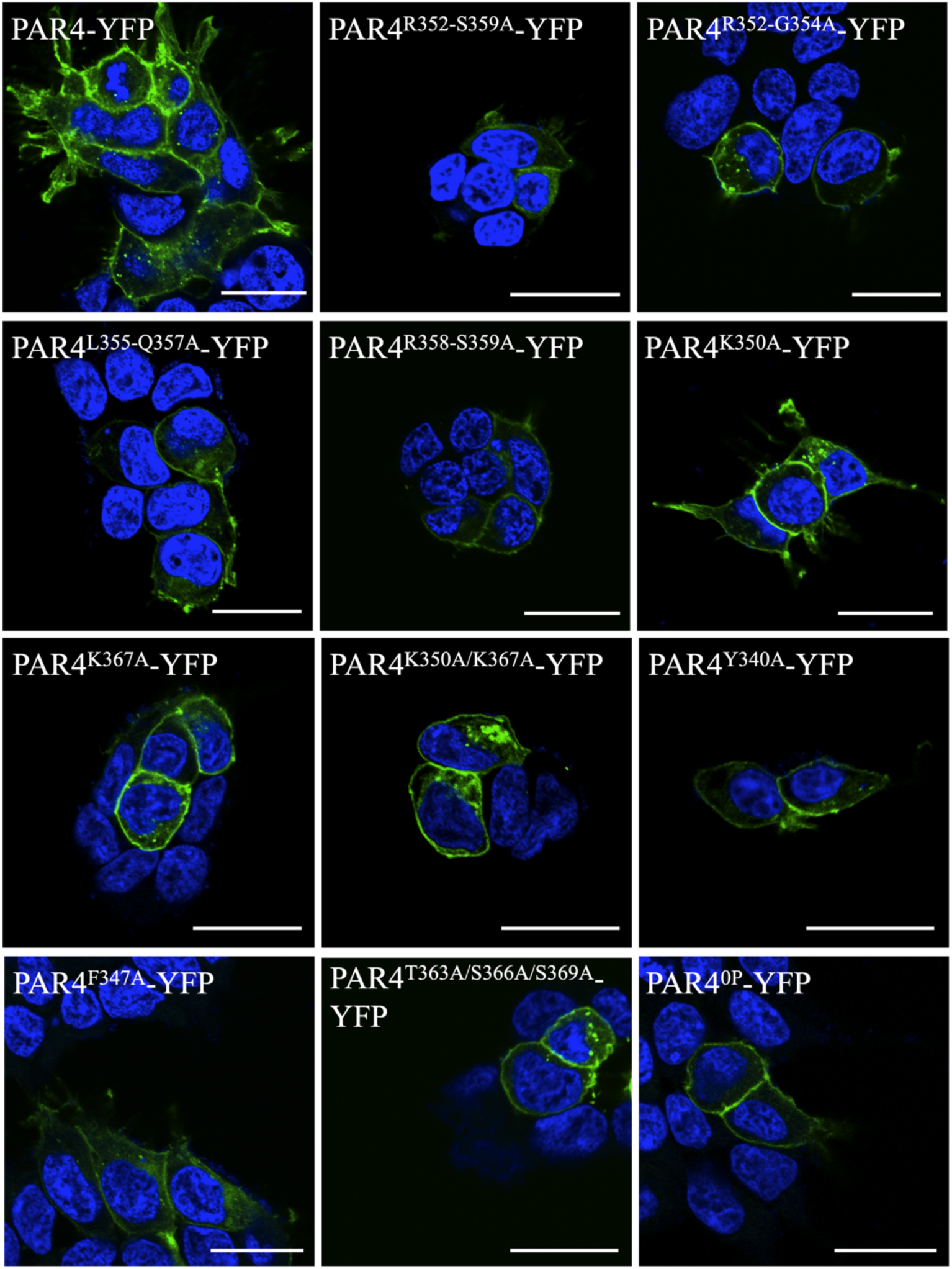
Confocal micrographs of PAR4-YFP and mutant receptor cellular expression. Confocal microscopy was employed to determine appropriate cell membrane expression of wild-type and mutant PAR4-YFP constructs. We observed membrane expression with wild-type and all mutant PAR4 receptor constructs. Scale bar is 20 µm. (*n = 3*)

**Supplementary Figure 2.**
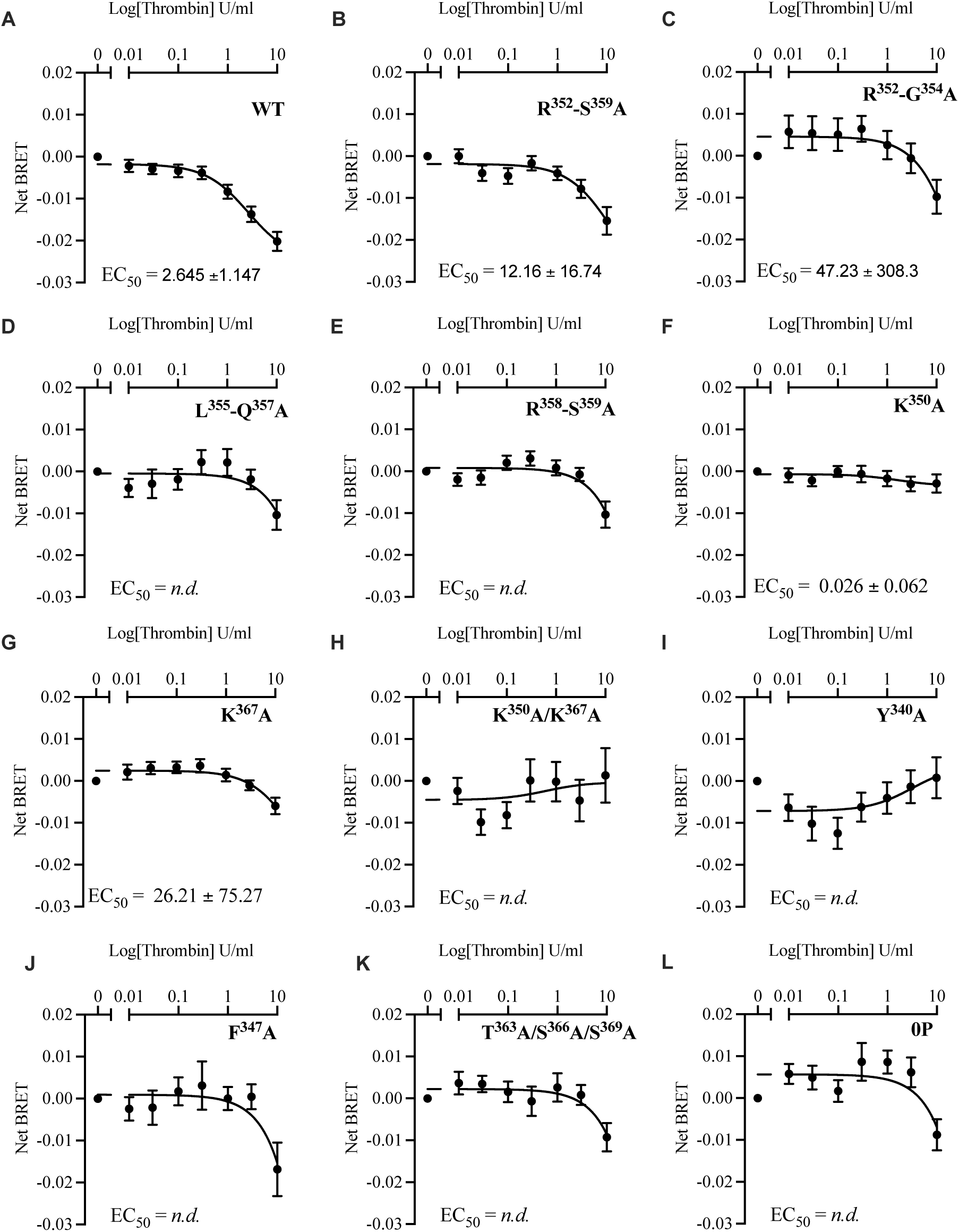
Gα_11_ activation (TRUPATH) by thrombin-stimulated wild-type (WT) and mutant PAR4 receptors. Nonlinear regression curve fits are shown (mean ± S.E.) for at least three independent experiments (*n = 3*) in PAR1-KO-HEK-293 cells.

**Supplementary Figure 3.**
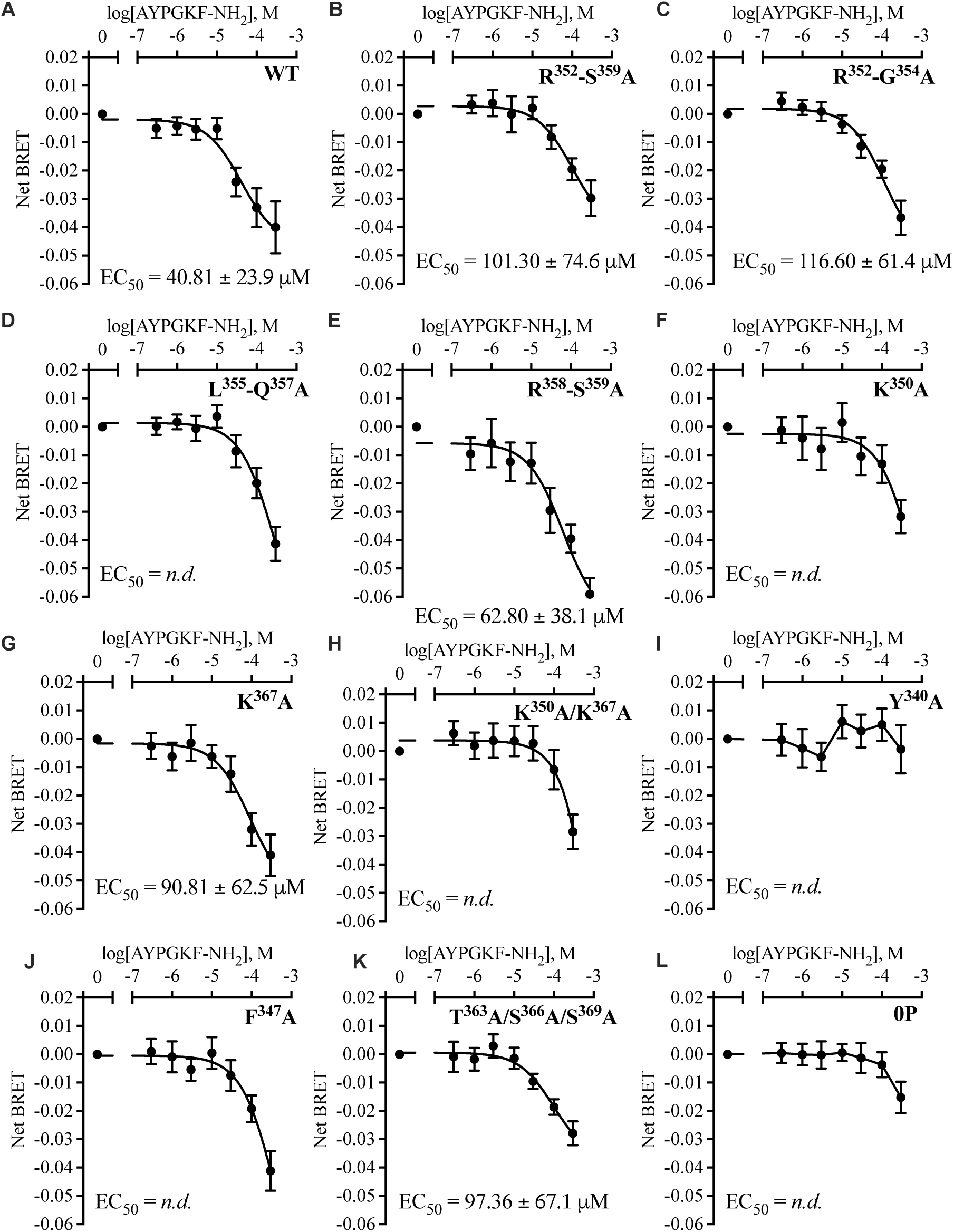
Gα_11_ activation (TRUPATH) by AYPGKF-NH_2_-stimulated wild-type (WT) and mutant PAR4 receptors. Nonlinear regression curve fits are shown (mean ± S.E.) for at least three independent experiments (*n = 3*) in PAR1-KO-HEK-293 cells.

